# Gut microbial taxa elevated by dietary sugar disrupt memory function

**DOI:** 10.1101/2020.06.16.153809

**Authors:** Emily E Noble, Christine A. Olson, Elizabeth Davis, Linda Tsan, Yen-Wei Chen, Ruth Schade, Clarissa Liu, Andrea Suarez, Roshonda B Jones, Claire de La Serre, Xia Yang, Elaine Y. Hsiao, Scott E Kanoski

## Abstract

Emerging evidence highlights a critical relationship between gut microbiota and neurocognitive development. Excessive consumption of sugar and other unhealthy dietary factors during early life developmental periods yields changes in the gut microbiome as well as neurocognitive impairments. However, it is unclear whether these two outcomes are functionally connected. Here we explore whether excessive early life consumption of added sugars negatively impacts memory function via the gut microbiome. Rats were given free access to a sugar-sweetened beverage (SSB) during the adolescent stage of development. Memory function and anxiety-like behavior were assessed during adulthood and gut bacterial and brain transcriptome analyses were conducted. Taxa-specific microbial enrichment experiments examined the functional relationship between sugar-induced microbiome changes and neurocognitive and brain transcriptome outcomes. Chronic early life sugar consumption impaired adult hippocampal-dependent memory function without affecting body weight or anxiety-like behavior. Adolescent SSB consumption during adolescence also altered the gut microbiome, including elevated abundance of two species in the genus *Parabacteroides* (*P. distasonis* and *P. johnsonii*) that were negatively correlated with hippocampal function. Transferred enrichment of these specific bacterial taxa in adolescent rats impaired hippocampal-dependent memory during adulthood. Hippocampus transcriptome analyses revealed that early life sugar consumption altered gene expression in intracellular kinase and synaptic neurotransmitter signaling pathways, whereas *Parabacteroides* microbial enrichment altered gene expression in pathways associated with metabolic function, neurodegenerative disease, and dopaminergic signaling. Collectively these results identify a role for microbiota “dysbiosis” in mediating the detrimental effects of early life unhealthy dietary factors on hippocampal-dependent memory function.

## Introduction

The gut microbiome has recently been implicated in modulating neurocognitive development and consequent functioning ^1-4^. Early life developmental periods represent critical windows for the impact of indigenous gut microbes on the brain, as evidenced by the reversal of behavioral and neurochemical abnormalities in germ free rodents when inoculated with conventional microbiota during early life, but not during adulthood ^5-7^. Dietary factors are a critical determinant of gut microbiota diversity and can alter gut bacterial communities, as evident from the microbial plasticity observed in response to pre- and probiotic treatment, as well as the “dysbiosis” resulting from consuming unhealthy, yet palatable foods that are associated with obesity and metabolic disorders (e.g., Western diet; foods high in saturated fatty acids and added sugar) ^8^. In addition to altering the gut microbiota, consumption of Western dietary factors yields long-lasting memory impairments, and these effects are more pronounced when consumed during early life developmental periods vs. during adulthood ^9-11^. Whether diet-induced changes in specific bacterial populations are functionally related to altered early life neurocognitive outcomes, however, is poorly understood.

The hippocampus, which is well known for its role in spatial and episodic memory and more recently for regulating learned and social aspects of food intake control ^12-17^, is particularly vulnerable to the deleterious effects of Western dietary factors ^18-20^. During the juvenile and adolescent stages of development, a time when the brain is rapidly developing, consumption of diets high in saturated fat and sugar ^21-23^ or sugar alone ^24-27^ impairs hippocampal function while in some cases preserving memory processes that do not rely on the hippocampus. While several putative underlying mechanisms have been investigated, the precise biological pathways linking dietary factors to neurocognitive dysfunction remain largely undetermined ^11^. Here we aimed to determine whether sugar-induced alterations in gut microbiota during early life are causally related to hippocampal-dependent memory impairments observed during adulthood.

## Methods and Materials

### Experimental Subjects

Juvenile male Sprague Dawley rats (Envigo; arrival post natal day (PN) 26-28; 50-70g) were housed individually in standard conditions with a 12:12 light/dark cycle. All rats had ad libitum access to water and Lab Diet 5001 (PMI Nutrition International, Brentwood, MO; 29.8 % kcal from protein, 13.4% kcal from fat, 56.7% kcal from carbohydrate), with modifications where noted. All experiments were performed in accordance with the approval of the Animal Care and Use Committee at the University of Southern California.

### Experiment 1

Twenty one juvenile male rats (PN 26-28) were divided into two groups with equal body weight and given ad libitum access to: 1) 11% weight-by-volume (w/v) solution containing monosaccharide ratio of 65% fructose and 35% glucose in reverse osmosis-filtered water (SUG; *n*=11) or 2) or an extra bottle of reverse osmosis-filtered water (CTL; *n*=10). This solution was chosen to model commonly consumed sugar-sweetened beverages in humans in terms of both caloric content and monosaccharide ratio^28^.

Additionally, all rats were given ad libitum access to water and standard rat chow. Food intake, solution intake and body weights were monitored thrice weekly except where prohibited due to behavioral testing. At PN 60, rats underwent Novel Object in Context (NOIC) testing, to measure hippocampal-dependent episodic contextual memory. At PN 67 rats underwent anxiety-like behavior testing in the Zero Maze, followed by body composition testing at PN 70 and an intraperitoneal glucose tolerance test (IP GTT) at PN 84. All behavioral procedures were run at the same time each day (4-6 hours into the light cycle). Fecal and cecal samples were collected prior to sacrifice at PN 104.

In a separate cohort of juvenile male rats (n=6/group) animals were treated as above, but on PN day 60 rats were tested in the Novel Object Recognition (NOR) and Open Field (OF) tasks, with two days in between tasks. Animals were sacrificed and tissue punches were collected from the dorsal hippocampus on PN day 65. Tissue punches were flash frozen in a beaker filled with isopentane and surrounded dry ice and then stored at -80°C until further analyses.

### Experiment 2

Twenty-three juvenile male rats (PN 26-28) were divided into two groups and received a gavage twice daily (12 hours apart) for 7 days (only one treatment was given on day 7) of either (1) saline (SAL; n=8), or (2) a cocktail of antibiotics consisting of Vancomycin (50 mg/kg), Neomycin (100 mg/kg), and Metronidazole (100 mg/kg) along with supplementation with 1 mg/mL of ampicillin in their drinking water (ABX; n=15), which is a protocol modified from ^29^. Animals were housed in fresh, sterile cages on Day 3 of the antibiotic or saline treatment, and again switched to fresh sterile cages on Day 7 after the final gavage. All animals were maintained on sterile, autoclaved water and chow for the remainder of the experiment. Rats in the ABX group were given water instead of ampicillin solution on Day 7. Animals in the ABX group were further subdivided to receive either gavage of a 1:1 ratio of *Parabacteroides distasonis* and *Parabacteroides johnsonii* (PARA; n=8) or saline (SAL; n=7) thirty six hours after the last ABX treatment. To minimize potential contamination, rats were handled minimally for 14 days. Cage changes occurred once weekly at which time animals and food were weighed. Experimenters wore fresh, sterile PPE and weigh boxes were cleaned with sterilizing solution in between each cage change. On PN 50 rats were tested in NOIC, on PN 60 rats were tested in NOR, on PN 62 rats were tested in the Zero Maze, followed by Open Field on PN 64. On PN 73 rats were given an IP GTT, and on PN 76 body composition was tested. Rats were sacrificed at PN 83 and dorsal hippocampus tissue punches and cecal samples were collected. Tissue punches were flash frozen in a beaker filled with isopentane and surrounded dry ice and cecal samples were placed in microcentrifuge tubes embedded in dry ice. Samples were subsequently stored at -80°C until further analyses.

### IP glucose tolerance test (IP GTT)

Animals were food restricted 24 hours prior to IP GTT. Immediately prior to the test, baseline blood glucose readings were obtained from tail tip and recorded by a blood glucose meter (One touch Ultra2, LifeScan Inc., Milpitas, CA). Each animal was then intraperitoneally (IP) injected with dextrose solution (0.923g/ml by body weight) and tail tip blood glucose readings were obtained at 30, 60, 90, and 120 min after IP injections, as previously described ^24^.

### Zero Maze

The Zero Maze is an elevated circular track (63.5 cm fall height, 116.8cm outside diameter), divided into four equal length sections. Two sections were open with 3 cm high curbs, whereas the 2 other closed sections contained 17.5 cm high walls. Animals are placed in the maze facing the open section of the track in a room with ambient lighting for 5 min while the experimenter watches the animal from a monitor outside of the room. The experimenter records the total time spent in the open sections (defined as the head and front two paws in open arms), and the number of crosses into the open sections from the closed sections.

### Novel object in context task (NOIC)

NOIC measures episodic contextual memory based on the capacity for an animal to identify which of two familiar objects it has never seen before in a specific context. Procedures were adapted from prior reports ^30,31^. Briefly, rats are habituated to two distinct contexts on subsequent days (with the habituation order counterbalanced by group) for 5-min sessions: Context 1 is a semi-transparent box (15in W x 24in L x 12in H) with orange stripes and Context 2 is a grey opaque box (17in W x 17in L x 16in H) (Context identify assignments counterbalanced by group), each context is in a separate dimly lit room, which is obtained using two desk lamps pointed toward the floor. Day 1 of NOIC begins with each animal being placed in Context 1 containing two distinct similarly sized objects placed in opposite corners: a 500ml jar filled with blue water (Object A) and a square glass container (Object B) (Object assignments and placement counterbalanced by group). On day 2 of NOIC, animals are placed in Context 2 with duplicates of one of the objects. On NOIC day 3, rats are placed in Context 2 with Objects A and Object B. One of these objects is not novel to the rat, but its placement in Context 2 is novel. All sessions are 5 minutes long and are video recorded. Each time the rat is placed in one of the contexts, it is placed with its head facing away from both objects. The time spent investigating each object is recorded from the video recordings by an experimenter who is blinded to the treatment groups. Exploration is defined as sniffing or touching the object with the nose or forepaws. The task is scored by calculating the time spent exploring the Novel Object to the context divided by the time spent exploring both Objects A and B combined, which is the novelty or “discrimination index”. Rats with an intact hippocampus will preferentially investigate the object that is novel to Context 2, given that this object is a familiar object yet is now presented in a novel context, whereas hippocampal inactivation impairs the preferential investigation of the object novel to Context 2 ^30^.

### Novel Object Recognition

The apparatus used for NOR is a grey opaque box (17in W x 17in L x 16in H) placed in a dimly lit room, which is obtained using two desk lamps pointed toward the floor. Procedures are adapted from ^32^. Rats are habituated to the empty arena and conditions for 10 minutes on the day prior to testing. The novel object and the side on which the novel object is placed is counterbalanced by group. The test begins with a 5-minute familiarization phase, where rats are placed in the center of the arena, facing away from the objects, with two identical copies of the same object to explore. The objects were either two identical cans or two identical bottles, counterbalanced by treatment group. The objects were chosen based on preliminary studies which determined that they are equally preferred by Sprague Dawley rats. Animals are then removed from the arena and placed in the home cage for 5 minutes. The arena and objects are cleaned with 10% ethanol solution, and one of the objects in the arena is replaced with a different one (either the can or bottle, whichever the animal has not previously seen, i.e., the “novel object”). Animals are again placed in the center of the arena and allowed to explore for 3 minutes. Time spent exploring the objects is recorded via video recording and analyzed using Any-maze activity tracking software (Stoelting Co., Wood Dale, IL).

### Open Field

Open field measures general activity level and also anxiety-like behavior in the rat. A large gray bin, 60 cm (L) X 56 CM (W) is placed under diffuse even lighting (30 lux). A center zone is identified and marked in the bin (19 cm L X 17.5 cm W). A video camera is placed directly overhead and animals are tracked using AnyMaze Software (Stoelting Co., Wood Dale, IL). Animals are placed in the center of the box facing the back wall and allowed to explore the arena for 10 min while the experimenter watches from a monitor in an adjacent room. The apparatus is cleaned with 10% ethanol after each rat is tested.

### Body Composition

Body composition (body fat, lean mass) was measured using LF90 time domain nuclear magnetic resonance (Bruker NMR minispec LF 90II, Bruker Daltonics, Inc.).

### Bacterial transfer

*Parabacteroides distasonis* (ATCC 8503) was cultured under anaerobic conditions at 37C in Reinforced Clostridial Medium (RCM, BD Biosciences). *Parabacteroides johnsonii* (DSM 18315) was grown in anaerobic conditions in PYG medium (modified, DSM medium 104). Cultures were authenticated by full-length 16S rRNA gene sequencing. For bacterial enrichment, 10^9^ colony-forming units of both *P. distasonis* and *P. johnsonii* were suspended in 500 µL pre-reduced PBS and orally gavaged into antibiotic-treated rats. When co-administered, a ratio of 1:1 was used for *P. distasonis* and *P. johnsonii*.

### Gut microbiota DNA extraction and 16s rRNA gene sequencing in sugar-fed and control rats

All samples were extracted and sequenced according to the guidelines and procedures established by the Earth Microbiome Project ^33^. DNA was extracted from fecal and cecal samples using the MO BIO PowerSoil DNA extraction kit. PCR targeting the V4 region of the 16S rRNA bacterial gene was performed with the 515F/806R primers, utilizing the protocol described in Caporaso et al.^34^. Amplicons were barcoded and pooled in equal concentrations for sequencing. The amplicon pool was purified with the MO BIO UltraClean PCR Clean-up kit and sequenced by the 2 x 150bp MiSeq platform at the Institute for Genomic Medicine at UCSD. All sequences were deposited in Qiita Study 11255 as raw FASTQ files. Sequences were demultiplexed using Qiime-1 based “split libraries” with the forward reads only dropping. Demultiplexed sequences were then trimmed evenly to 100 bp and 150 bp to enable comparison to other studies for meta-analyses. Trimmed sequences were matched to known OTUs at 97% identity.

### Gut microbiota DNA extraction and 16S rRNA gene sequencing for *Parabacteroides*-enriched and control rats

Total bacterial genomic DNA was extracted from rat fecal samples (0.25 g) using the Qiagen DNeasy PowerSoil Kit. The library was prepared following methods from (Caporaso et al., 2011). The V4 region (515F-806R) of the 16S rDNA gene was PCR amplified using individually barcoded universal primers and 30 ng of the extracted genomic DNA. The conditions for PCR were as follows: 94°C for 3 min to denature the DNA, with 35 cycles at 94°C for 45 s, 50°C for 60 s, and 72°C for 90 s, with a final extension of 10 min at 72°C. The PCR reaction was set up in triplicate, and the PCR products were purified using the Qiaquick PCR purification kit (QIAGEN). The purified PCR product was pooled in equal molar concentrations quantified by nanodrop and sequenced by Laragen, Inc. using the Illumina MiSeq platform and 2 x 250bp reagent kit for paired-end sequencing. Amplicon sequence variants (ASVs) were chosen after denoising with the Deblur pipeline. Taxonomy assignment and rarefaction were performed using QIIME2-2019.10.

### Hippocampal RNA extraction and sequencing

Hippocampi from rats treated with or without sugar or *Parabacteroides* were subject to RNA-seq analysis. Total RNA was extracted according to manufacturer’s instructions using RNeasy Lipid Tissue Mini Kit (Qiagen, Hilden, Germany). Total RNA was checked for degradation in a Bioanalyzer 2100 (Agilent, Santa Clara, CA, USA). Quality was very high for all samples, and libraries were prepared from 1 ug of total RNA using a NuGen Universal Plus mRNA-seq Library Prep Kit (Tecan Genomics Inc. Redwood City, CA). Final library products were quantified using the Qubit 2.0 Fluorometer (Thermo Fisher Scientific Inc., Waltham, MA, USA), and the fragment size distribution was determined with the Bioanalyzer 2100. The libraries were then pooled equimolarly, and the final pool was quantified via qPCR using the Kapa Biosystems Library Quantification Kit, according to manufacturer’s instructions. The pool was sequenced in an Illumina NextSeq 550 platform (Illumina, San Diego, CA, USA), in Single-Read 75 cycles format, obtaining about 25 million reads per sample. The preparation of the libraries and the sequencing was performed at the USC Genome Core (http://uscgenomecore.usc.edu/)

### RNA-seq quality control

Data quality checks were performed using the FastQC tool (http://www.bioinformatics.babraham.ac.uk/projects/fastqc) and low quality reads were trimmed with Trim_Galore (http://www.bioinformatics.babraham.ac.uk/projects/trim_galore/). RNA-seq reads passing quality control were mapped to *Rattus novegicus* transcriptome (Rnor6) and quantified with Salmon ^35^. Salmon directly mapped RNA-seq reads to Rat transcriptome and quantified transcript counts. Txiimport ^36^ were used to convert transcript counts into gene counts. Potential sample outliers were detected by principle component analysis (PCA) and one control and one treatment sample from the *Parabacteroides* experiment were deemed outliers (Figure S7A, B) and removed.

### Identification of differentially expressed genes (DEGs)

DESeq2^37^ were used to conduct differential gene expression analysis between sugar treatment and the corresponding controls, or between *Parabacteroides* treatment and the corresponding controls. Low-abundance genes were filtered out and only those having a mean raw count > 1 in more than 50% of the samples were included. Differentially expressed genes were detected by DESeq2 with default settings. Significant DEGs were defined as Benjamini-Hochberg (BH) adjusted false discovery rate (FDR) < 0.05. For heatmap visualization, genes were normalized with variance stabilization transformation implemented in DESeq2, followed by calculating a z-score for each gene.

### Pathway analyses of DEGs

For the pathway analyses, DEGs at an unadjusted p-value < 0.01 were used. Pathway enrichment analysis were conducted using enrichr^38^ by intersecting each signature with pathways or gene sets from KEGG^39^, gene ontology biological pathways (GOBP), Cellular Component (GOCP), Molecular Function (GOMF)^40^ and Wikipathways^41^. Pathways at FDR < 0.05 were considered significant. Unless otherwise specified, R 3.5.2 was used for the analysis mentioned in the RNA sequencing section.

### Additional statistical methods

Data are presented as means ± SEM. For analytic comparisons of body weight, total food intake, and chow intake, groups were compared using repeated measures ANOVA in Prism software (GraphPad Inc., version 8.0). Taxonomic comparisons from 16S rRNA sequencing analysis were analyzed by analysis of composition of microbiomes (ANCOM). When significant differences were detected, Sidak post-hoc test for multiple comparisons was used. Area under the curve (AUC) for the IP GTT testing was also calculated using Prism. All other statistical analyses were performed using Student’s two-tailed unpaired t tests in excel software (Microsoft Inc., version 15.26). Normality was confirmed prior to the utilization of parametric testing. For all analyses, statistical significance was set at *P*<0.05.

## Results

### Early-life sugar consumption impairs hippocampal-dependent memory function

Results from the Novel Object in Context (NOIC) task, which measures hippocampal-dependent episodic contextual memory function ^31^, reveal that while there were no differences in total exploration time of the combined objects on days 1 or 3 of the task (Figure 1A, B), animals fed sugar solutions in early life beginning at PN 28 had a reduced capacity to discriminate an object that was novel to a specific context when animals were tested during adulthood (PN 60), indicating impaired hippocampal function (Figure 1C). Conversely, animals fed sugar solutions in early life performed similarly to those in the control group when tested in the novel object recognition task (NOR) (Figure 1D), which tests object recognition memory independent of context. Notably, when performed using the current methods with a short duration between the familiarization phase and the test phase, NOR not hippocampal-dependent but instead is primarily dependent on the perirhinal cortex ^31,42-44^. These data suggest that early life dietary sugar consumption impairs performance in hippocampal-dependent contextual-based recognition memory without affecting performance in perirhinal cortex-dependent recognition memory independent of context ^24^.

**Figure 1:**
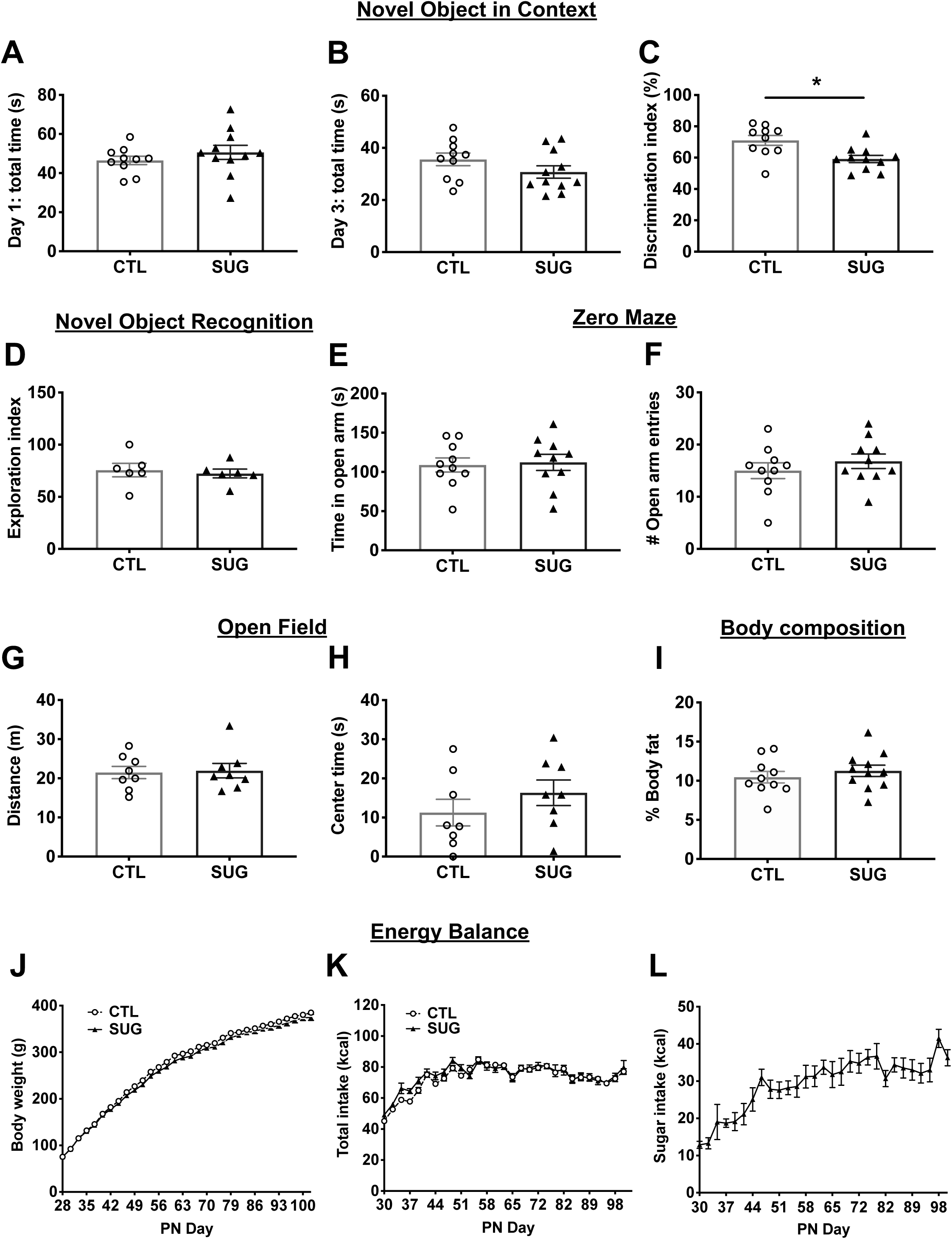
Early-life sugar consumption negatively impacts hippocampal-dependent memory function. (A, B) Early life sugar consumption had no effect on total exploration time on days 1 (familiarization) or day 3 (test day) of the Novel Object in Context (NOIC) task. (C) The discrimination index was significantly reduced by early life sugar consumption, indicating impaired hippocampal function (P<.05, n=10,11; two-tailed, type 2 Student’s T-test). (D) There were no differences in exploration index in the Novel Object Recognition (NOR task) (n=6; two-tailed, type 2 Student’s T-test). (E, F) There were no differences in time spent in the open arm or the number of entries into the open arm in the Zero Maze task for anxiety-like behavior (n=10; two-tailed, type 2 Student’s t-test). (G, H) There were no differences in distance travelled or time spent in the center arena in the Open Field task (n=8; two-tailed, type 2 Student’s T-test). (I) There were no differences in body fat % during adulthood between rats fed early life sugar and controls (n=10,11; two-tailed, type 2 Student’s T-test). (J-K) Body weights and total energy intake did not differ between the groups (n=10,11; two-way repeated measures ANOVA), despite (L) increased kcal consumption from sugar sweetened beverages in the sugar group. CTL=control, SUG= sugar, PN= post-natal day; data shown as mean ± SEM.

Elevated anxiety-like behavior and altered general activity levels may influence novelty exploration independent of memory effects and may therefore confound the interpretation of behavioral results. Thus, we next tested whether early life sugar consumption affects anxiety-like behavior using two different tasks designed to measure anxiety-like behavior in the rat: the elevated zero maze and the open field task, that latter of which also assesses levels of general activity ^45^. Early life sugar consumption had no effect on time spent in the open area or in the number of open area entries in the zero maze (Figure 1E, F). Similarly, early life sugar had no effect on distance travelled or time spent in the center zone in the open field task (Figure 1G, H). Together these data suggest that habitual early life sugar consumption did not increase anxiety-like behavior or general activity levels in the rats.

### Early life sugar consumption impairs glucose tolerance without affecting total caloric intake, body weight, or adiposity

Given that excessive sugar consumption is associated with weight gain and metabolic deficits ^46^, we tested whether access to a sugar solution during the adolescent phase of development would affect food intake, body weight gain, adiposity, and glucose tolerance in the rat. Early life sugar consumption had no effect on body composition during adulthood (Figure 1I, Figure S1 A, B). Early life sugar consumption also had no effect on body weight or total kcal intake (Figure 1J, K), which is in agreement with previous findings ^24,27,47^. Animals steadily increased their intake of the 11% sugar solution throughout the study (Figure 1L) but compensated for the calories consumed in the sugar solutions by reducing their intake of dietary chow (Figure S1 C). However, animals that were fed sugar solutions during adolescence showed impaired peripheral glucose metabolism in an intraperitoneal glucose tolerance test (IP GTT) (Figure S1D).

### Gut microbiota are impacted by early life sugar consumption

Principal component analyses of 16s rRNA gene sequencing data of fecal samples revealed a separation between the fecal microbiota of rats fed early life sugar and controls (Figure 2A). Results from LEfSe analysis identified differentially abundant bacterial taxa in fecal samples that were elevated by sugar consumption. These include the family *Clostridiaceae* and the genus *02d06* within *Clostridiaceae*, the family *Mogibacteriaceae*, the family *Enterobacteriaceae*, the order *Enterobacteriales*, the class of *Gammaproteobacteria*, and the genus *Parabacteroides* within the family *Porphyromonadaceae* (Figure 2B,C). In addition to an elevated % relative abundance of the genus *Parabacteroides* in animals fed early life sugar (Figure 2D), log transformed counts of the *Parabacteroides* negatively correlated with performance scores in the NOIC memory task (Figure 2E). Of the additional bacterial populations significantly affected by sugar treatment, regression analyses did not identify any other genera as being significantly correlated to NOIC memory performance. Within *Parabacteroides*, levels of two operational taxonomic units (OTUs) that were elevated by sugar negatively correlated with performance in the NOIC task, identified as taxonomically related to *P. johnsonii* and *P. distasonis* (Figure 2F, G). The significant negative correlation between NOIC performance and *Parabacteroides* was also present within each of the diet groups alone, but when separated out by diet group only *P. distasonis* showed a significant negative correlation for each diet group (P<.05), whereas *P. johnsonii* showed a non-significant trend in both the control and sugar groups (P=.06, and P=.08, respectively; Figure S2A-C). Abundance of other bacterial populations that were affected by sugar consumption were not significantly related to memory task performance.

**Figure 2:**
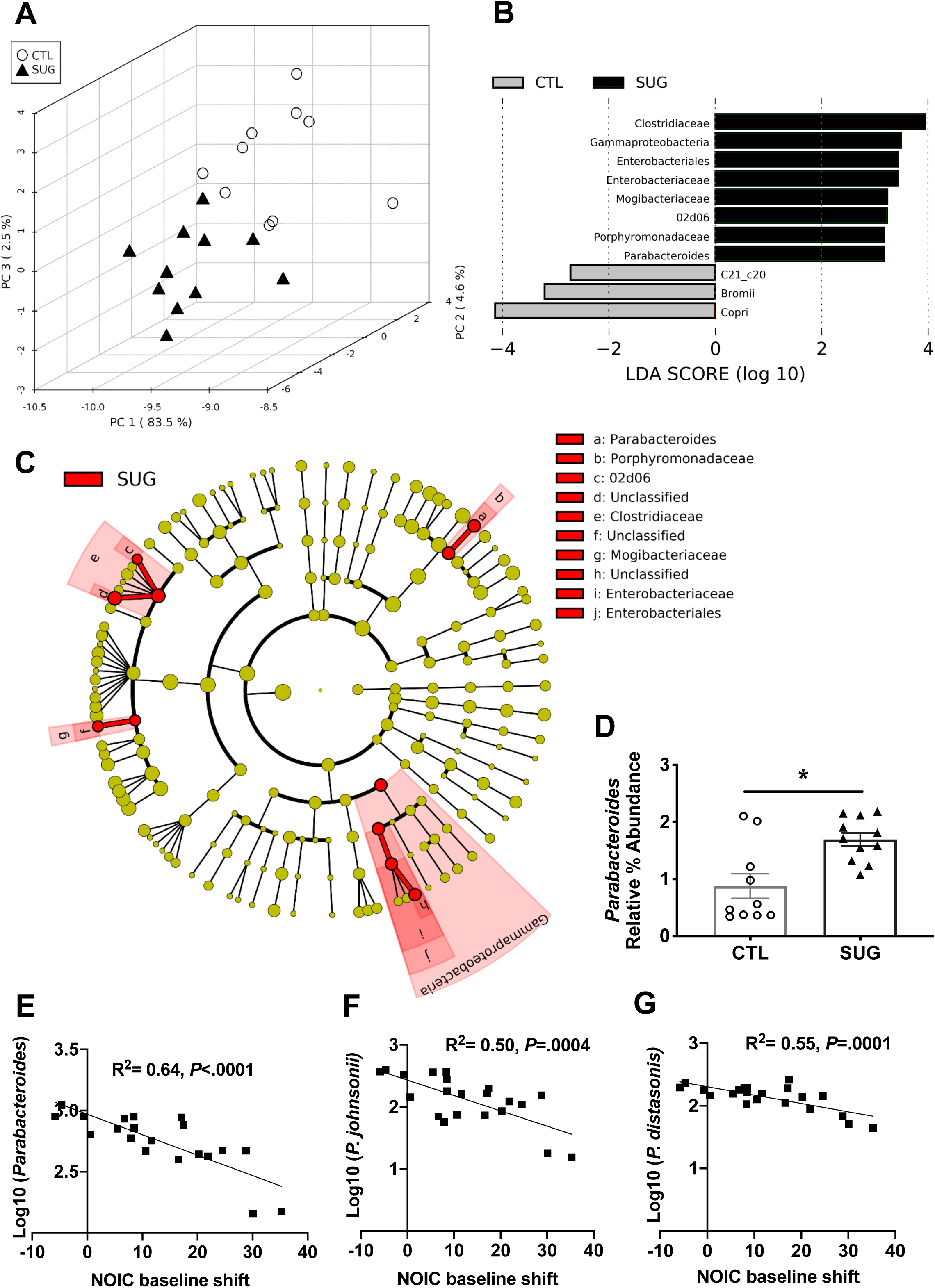
Effect of adolescent sugar consumption on the gut microbiome in rats. (A) Principal component analysis showing separation between fecal microbiota of rats fed early life sugar or controls (n=11, 10; dark triangles= sugar, open circles= control). (B) Results from LEfSe analysis showing Linear Discriminate Analysis (LDA) scores for microbiome analysis of fecal samples of rats fed early life sugar or controls. (C) A cladogram representing the results from the LEfSe analysis with class as the outer most taxonomic level and species at the inner most level. Taxa in red are elevated in the sugar group. (D) Relative % abundance of fecal *Parabacteroides* are significantly elevated in rats fed early life sugar (P<.05; n=11, 10, two-tailed, type 2 Student’s T-test). (E) Linear regression of log normalized fecal *Parabacteroides* counts against shift from baseline performance scores in the novel object in context task (NOIC) across all groups tested (n=21). (E, F) Linear regression of the most abundant fecal *Parabacteroides* species against shift from baseline performance scores in NOIC across all groups tested (n=21). *P<0.05; data shown as mean ± SEM.

There was a similar separation between groups in bacteria analyzed from cecal samples (Figure S3A). LEfSe results from cecal samples show elevated *Bacilli*,

*Actinobacteria*, *Erysipelotrichia*, and *Gammaproteobacteria* in rats fed early life sugar, and elevated *Clostridia* in the controls (Figure S3B-C). Abundances at the different taxonomic levels in fecal and cecal samples are shown in (Figure S4, S5). Regression analyses did not identify these altered cecal bacterial populations as being significantly correlated to NOIC memory performance.

### Early life *Parabacteroides* enrichment impairs memory function

To determine whether neurocognitive outcomes due to early life sugar consumption could be attributable to elevated levels of *Parabacteroides* in the gut, we experimentally enriched the gut microbiota of naïve juvenile rats with two *Parabacteroides* species that exhibited high 16S rRNA sequencing alignment with OTUs that were increased by sugar consumption and were negatively correlated with behavioral outcomes in rats fed early life sugar. *P. johnsonii* and *P. distasoni* species were cultured individually under anaerobic conditions and transferred to a group of antibiotic-treated young rats in a 1:1 ratio via oral gavage using the experimental design described in Methods and outlined in Figure 3A, and from ^29^. To confirm *Parabacteroides* enrichment, 16SrRNA sequencing was performed on rat fecal samples for SAL-SAL, ABX-SAL, and ABX-PARA groups. Alpha diversity was analyzed using observed operational taxonomic units (OTUs) (Figure 3B), where both ABX-SAL and ABX-PARA fecal samples have significantly reduced alpha diversity when compared with SAL-SAL fecal samples, suggesting that antibiotic treatment reduces microbiome alpha diversity. Further, either treatment with antibiotics alone or antibiotics followed by *Parabacteroides* significantly alters microbiota composition relative to the SAL-SAL group (Figure 3C). Taxonomic comparisons from 16S rRNA sequencing analysis were analyzed by analysis of composition of microbiomes (ANCOM). Differential abundance on relative abundance at the species level (Figure 3D) was tested across samples hypothesis-free. Significant taxa at the species level were corrected for using false-discovery rate (FDR)-corrected *P*-values to calculate W in ANCOM. Comparing all groups resulted in the highest W value of 144 for the *Parabacteroides* genus, which was enriched in ABX-PARA fecal samples after bacterial gavage with an average relative abundance of 55.65% (Figure 3E). This confirms successful *Parabacteroides* enrichment for ABX-PARA rats post-gavage when compared to either ABX-SAL (average relative abundance of 5.47%) or ABX-SAL rats (average relative abundance of 0.26%).

**Figure 3:**
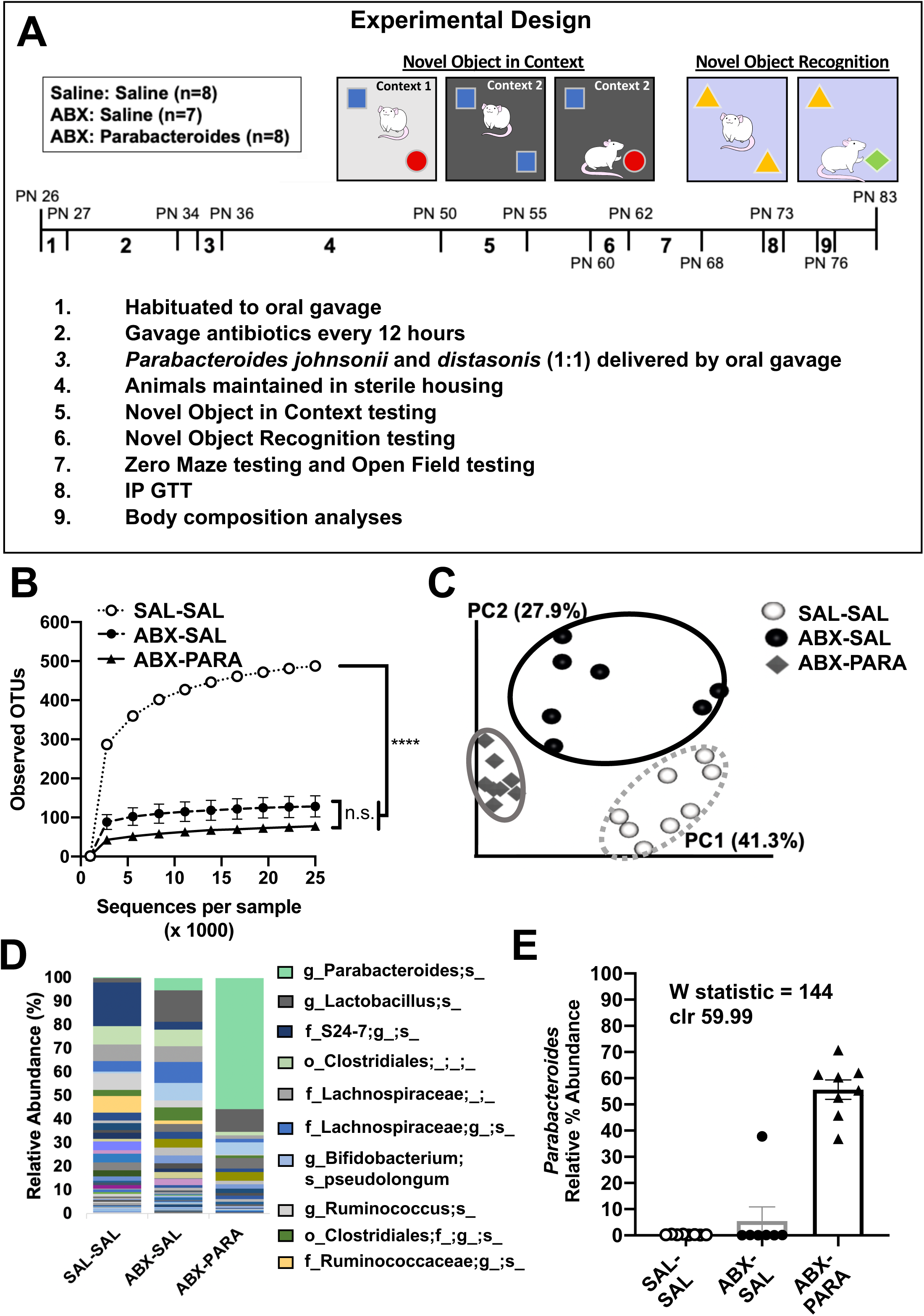
Intestinal *Parabacteroides* is enriched by antibiotic treatment and oral gavage of *P. distasonis* and *P. johnsonii*. A) Schematic showing the timeline for the experimental design of the *Parabacteroides* transfer experiment. B) Alpha diversity based on 16S rRNA gene profiling of fecal matter (n=7-8) represented by observed operational taxonomic units (OTUs) for a given number of sample sequences. C) Principal coordinates analysis of weighted UniFrac distance based on 16S rRNA gene profiling of feces for SAL-SAL, ABX-SAL, and ABX-PARA enriched rats (n=7-8). D) Average taxonomic distributions of bacteria from 16S rRNA gene sequencing data of feces for SAL-SAL, ABX-SAL, and ABX-PARA enriched animals (n=7-8). E) Relative abundances of *Parabacteroides* in fecal microbiota for SAL-SAL, ABX-SAL, and ABX-PARA enriched animals (n=7-8) (ANCOM). PN= post-natal day, IP GTT= intraperitoneal glucose tolerance test. Data are presented as mean ± S.E.M. * *p* < 0.05, \*\**p* < 0.01, \*\*\**p* < 0.001. n.s.=not statistically significant. SAL-SAL= rats treated with saline, ABX-SAL=rats treated with antibiotics followed by sterile saline gavage, ABX-PARA= rats treated with antibiotics followed by a 1:1 gavage of *Parabacteroides distasonis* and *Parabacteroides johnsonii*.

All rats treated with antibiotics showed a reduction in food intake and body weight during the initial stages of antibiotic treatment, however, there were no differences in body weight between the two groups of antibiotic treated animals by PN50, at the time of behavioral testing (Figure S6A-C). Similar to a recent report ^48^, *Parabacteroides* enrichment in the present study impacted body weight at later time points. Animals who received *P. johnsonii* and *P. distasonis* treatment showed reduced body weight 40 days after the transfer, with significantly lower lean mass (Figure S6D-F). There were no differences in percent body fat between groups, nor were there significant group differences in glucose metabolism in the IPGTT (Figure S6 G). Importantly, the body weights in the ABX-PARA group did not significantly differ from the ABX-SAL control group at the time of behavioral testing.

Results from the hippocampal-dependent NOIC memory task showed that while there were no differences in total exploration time of the combined objects on days 1 or 3 of the task, indicating similar exploratory behavior, animals enriched with *Parabacteroides* showed a significantly reduced discrimination index in the NOIC task compared with either control group (Figure 4A-C), indicating impaired performance in hippocampal-dependent memory function. When tested in the perirhinal cortex-dependent NOR task ^31^, animals enriched with *Parabacteroides* showed impaired object recognition memory compared with the antibiotic treated control group as indicated by a reduced novel object exploration index (Figure 4D). These findings show that unlike sugar-fed animals, *Parabacteroides* enrichment impaired perirhinal cortex-dependent memory processes in addition to hippocampal-dependent memory.

**Figure 4:**
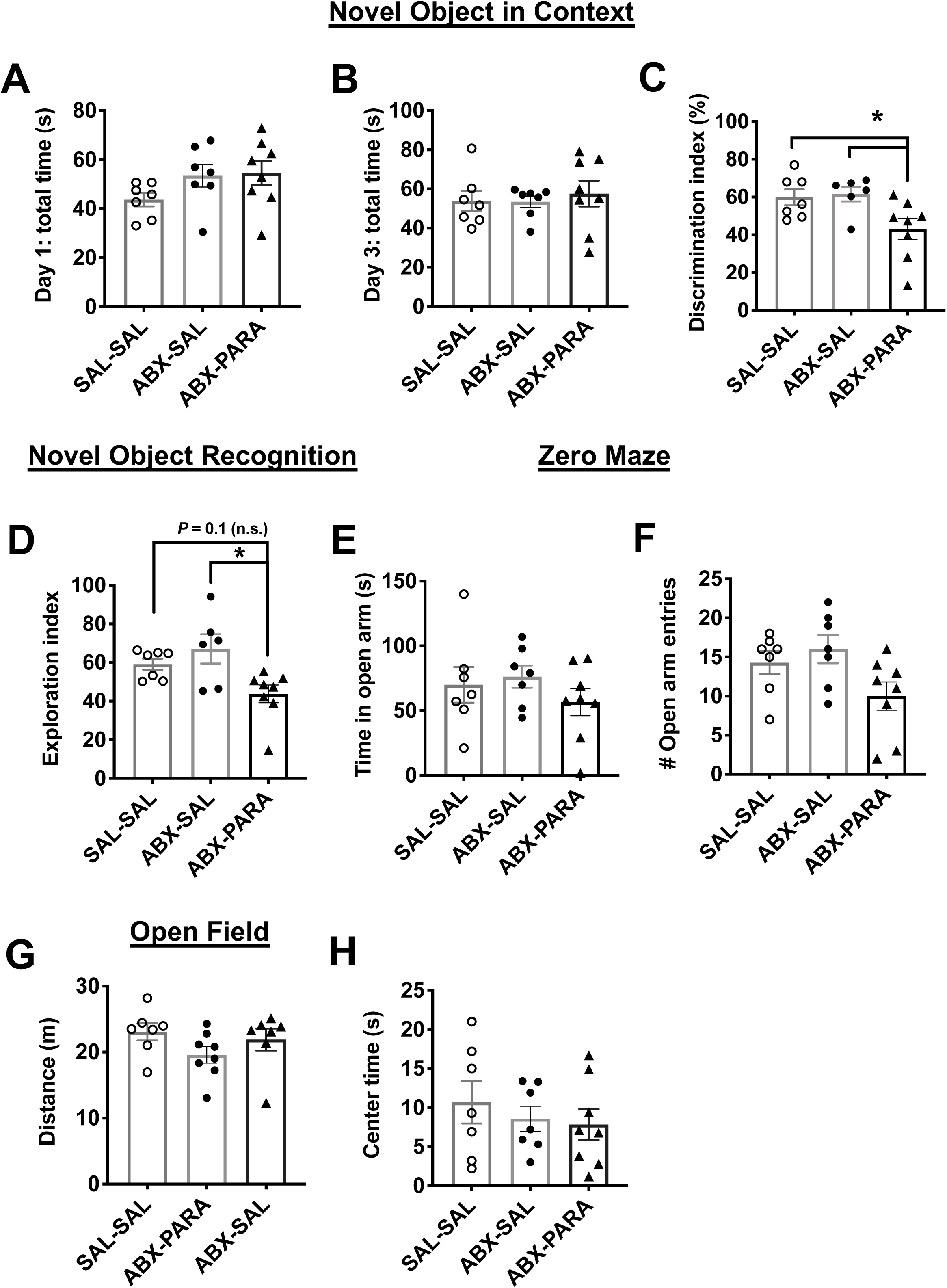
Early-life enrichment with *Parabacteroides* negatively impacts neurocognitive function. (A, B) Early-life enrichment with a 1:1 ratio of *P. johnsonii* and *P. distasonis* had no effect on total exploration time in the Novel Object in Context (NOIC) task. (C) Discrimination index was significantly reduced by enrichment with *P. johnsonii* and *P. distasonis*, indicating impaired hippocampal function (n=7,8; F _(2, 19)_ = 4.92; P< .05, one-way ANOVA with Tukey’s multiple comparison test). (D) There was a significant reduction in the exploration index in the Novel Object Recognition (NOR task), indicating impaired perirhinal cortex function (n=7,8; F _(2, 19)_ = 3.61; P< .05, one-way ANOVA with Tukey’s multiple comparison test). (E, F) There were no differences in time spent or number of entries into the open arm by animals with *P. johnsonii* and *P. distasonis* enrichment in the Zero Maze task for anxiety-like behavior (n=7,8; one-way ANOVA). (G, H) There were no differences in distance travelled or time spent in the center arena in the Open Field task (n=7,8; one-way ANOVA). SAL-SAL=saline-saline control, ABX-SAL= antibiotics-saline control, ABX-PARA= antibiotics-*P. johnsonii* and *P. distasonis* enriched, PN= post-natal day; data shown as mean ± SEM; * *P*<.05.

Results from the zero maze showed no differences in time spent in the open arms nor in the number of open arm entries for the *Parabacteroides*-enriched rats relative to controls (Figure 4E, F), indicating that the enrichment did not affect anxiety-like behavior. Similarly, there were no differences in distance travelled or time spent in the center arena in the open field test, which is a measure of both anxiety-like behavior and general activity in rodents (Figure 4G, H). Together these data suggest that *Parabacteroides* treatment negatively impacted both hippocampal-dependent perirhinal cortex-dependent memory function without significantly affecting general activity or anxiety-like behavior.

### Early life sugar consumption and *Parabacteroides* enrichment alter hippocampal gene expression profiles

To further investigate how sugar and *Parabacteroides* affect cognitive behaviors, we conducted transcriptome analysis of the hippocampus samples. Figure S7 (A, C) shows the results of principal component analysis revealing moderate separation based on RNA sequencing data from the dorsal hippocampus of rats fed sugar in early life compared with controls. Gene pathway enrichment analyses from RNA sequencing data revealed multiple pathways significantly affected by early life sugar consumption, including four pathways involved in neurotransmitter synaptic signaling: dopaminergic, glutamatergic, cholinergic, and serotonergic signaling pathways. Additionally, several gene pathways that also varied by sugar were those involved in kinase-mediated intracellular signalling: cGMP-PKG, RAS, cAMP, and MAPK signaling pathways (Figure 5A, Table S1).

**Figure 5:**
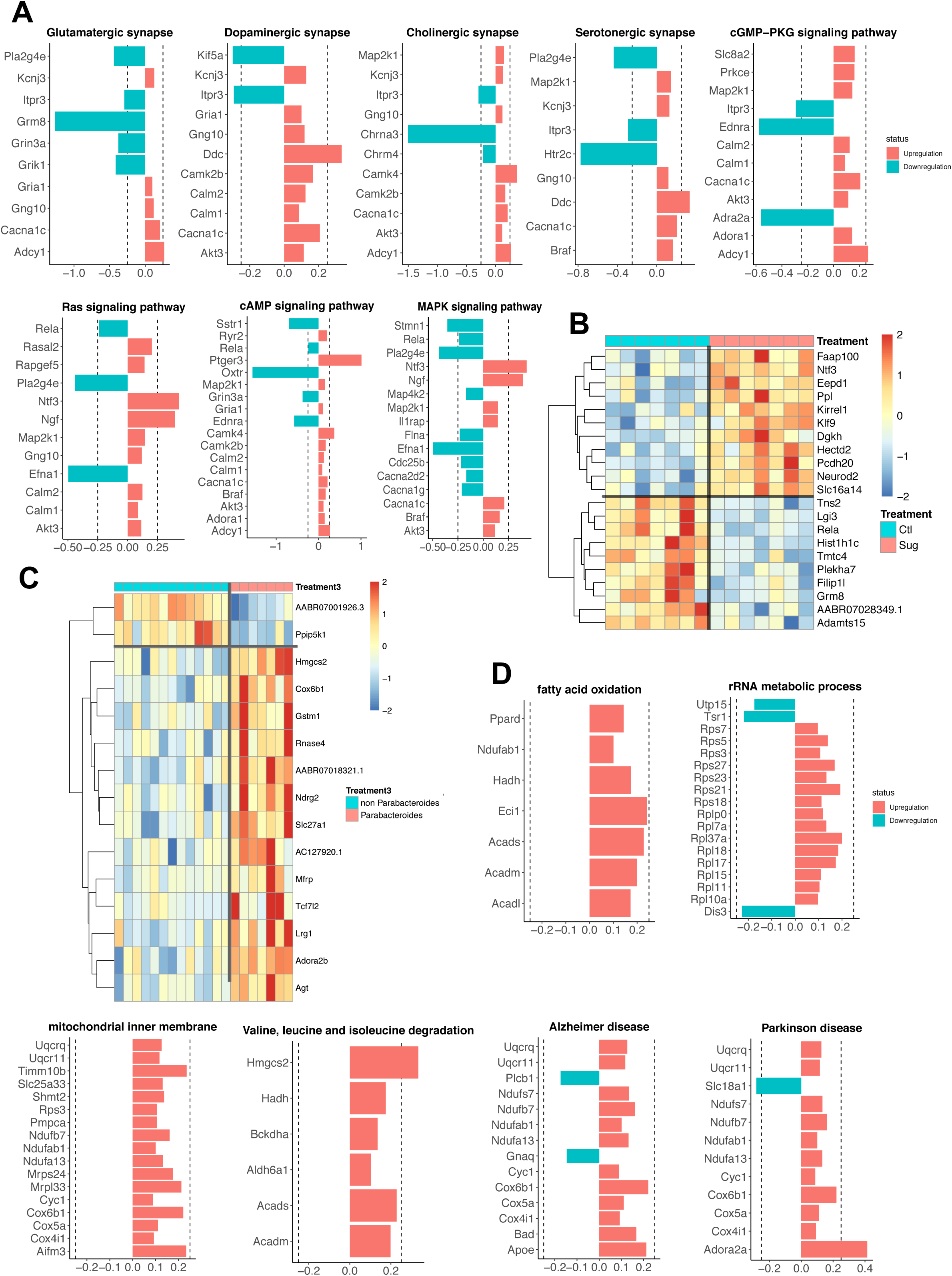
Effect of early life sugar or targeted *Parabacteroides* enrichment on hippocampal gene expression. (A) Pathway analyses for differentially expressed genes (DEGs) at a p-value < 0.01 in hippocampal tissue punches from rats fed early life sugar compared with controls. Upregulation by sugar is shown in red and downregulation by sugar in blue. (B) A heatmap depicting DEGs that survived the Benjamini-Hochberg corrected FDR of P< 0.05 in rats fed early life sugar compared with controls. Warmer colors (red) signify an increase in gene expression and cool colors (blue) a reduction in gene expression by treatment (CTL=control, SUG= early life sugar; n=7/group). (C) A heatmap depicting DEGs that survived the Benjamini-Hochberg corrected FDR of P< 0.05 in rats with early life *Parabacteroides* enrichment compared with combined control groups. Warmer colors (red) signify an increase in gene expression and cool colors (blue) a reduction in gene expression by treatment (n=7, 14). (D) Pathway analyses for differentially expressed genes (DEGs) at a P-value < 0.01 in rats enriched with *Parabacteroides* compared with combined controls. Upregulation by *Parabacteroides* transfer is shown in red and downregulation in blue. Dotted line indicates ±0.25 log2 fold change.

Analyses of individual genes across the entire transcriptome using a stringent false-discovery rate criterion further identified 21 genes that were differentially expressed in rats fed early life sugar compared with controls, with 11 genes elevated and 10 genes decreased in rats fed sugar compared to controls (Figure 5B). Among the genes impacted, several genes that regulate cell survival, migration, differentiation, and DNA repair were elevated by early life sugar access, including *Faap100*, which encodes an FA core complex member of the DNA damage response pathway ^49^, and *Eepd1*, which transcribes an endonuclease involved in repairing stalled DNA replication forks, stressed from DNA damage ^50^. Other genes associated with endoplasmic reticulum stress and synaptogenesis were also significantly increased by sugar consumption, including *Klf9, Dgkh, Neurod2, Ppl*, and *Kirrel1* ^51,52,53,54^.

Several genes were reduced by dietary sugar, including *Tns2*, which encodes tensin 2, important for cell migration ^55^, *RelA*, which encodes a NF/kB complex protein that regulates activity dependent neuronal function and synaptic plasticity ^56^, and *Grm8*, the gene for the metabotropic glutamate receptor 8 (mGluR8). Notably, reduced expression of mGluR8 receptor may contribute to the impaired neurocognitive functioning in animals fed sugar, as mGluR8 knockout mice show impaired hippocampal-dependent learning and memory ^57^.

Figure S7 (A-B, D) shows the results of principal component analysis of dorsal hippocampus RNA sequencing data indicating moderate separation between rats enriched with *Parabacteroides* and controls. Gene pathway analyses revealed that early life *Parabacteroides* treatment, similar to effects associated with sugar consumption, significantly altered the genetic signature of dopaminergic synaptic signaling pathways, though differentially expressed genes were commonly affected in opposite directions between the two experimental conditions (Figure S8). *Parabacteroides* treatment also impacted gene pathways associated with metabolic signaling. Specifically, pathways regulating fatty acid oxidation, rRNA metabolic processes, mitochondrial inner membrane, and valine, leucine, and isoleucine degradation were significantly affected by *Parabacteroides* enrichment. Other pathways that were influenced were those involved in neurodegenerative disorders, including Alzheimer’s disease and Parkinson’s disease, though most of the genes affected in these pathways were mitochondrial genes (Figure 5D, Table S2).

At the level of individual genes, dorsal hippocampal RNA sequencing data revealed that 15 genes were differentially expressed in rats enriched with *Parabacteroides* compared with controls, with 13 genes elevated and two genes decreased in the *Parabacteroides* group compared with controls (Figure 6C). Consistent with results from gene pathway analyses, several individual genes involved in metabolic processes were elevated by *Parabacteroides* enrichment, such as *Hmgcs2*, which is a mitochondrial regulator of ketogenesis and provides energy to the brain under metabolically taxing conditions or when glucose availability is low ^58^, and *Cox6b1*, a mitochondrial regulator of energy metabolism that improves hippocampal cellular viability following ischemia/reperfusion injury ^59^. *Parabacteroides* enrichment was also associated with incased expression of *Slc27A1* and *Mfrp*, which are each critical for the transport of fatty acids into the brain across capillary endothelial cells ^60,61^.

## Discussion

Dietary factors are a key source of gut microbiome diversity ^29,47,62-64^ and emerging evidence indicates that diet-induced alterations in the gut microbiota may be linked with altered neurocognitive development ^29,64-66^. Our results identify species within the genus *Parabacteroides* that are elevated by habitual early life consumption of dietary sugar and are negatively associated with hippocampal-dependent memory performance. Further, targeted microbiota enrichment of *Parabacteroides* perturbed both hippocampal- and perirhinal cortex-dependent memory performance. These findings are consistent with previous literature in showing that early life consumption of Western dietary factors impair neurocognitive outcomes ^10,11^, and further suggest that altered gut bacteria due to excessive early life sugar consumption may functionally link dietary patterns with cognitive impairment.

Our previous data show that rats are not susceptible to habitual sugar consumption-induced learning and memory impairments when 11% sugar solutions are consumed ad libitum during adulthood, in contrast to effects observed in the present and previous study in which the sugar is consumed during early life development ^24^. It is possible that habitual sugar consumption differentially affects the gut microbiome when consumed during adolescence vs. adulthood. However, a recent report showed that adult consumption of a high fructose diet (35% kcal from fructose) promotes gut microbial “dysbiosis” and neuroinflammation and cell death in the hippocampus, yet without impacting cognitive function ^67^, suggesting that perhaps neurocognitive function is more susceptible to gut microbiota influences during early life than during adulthood. Indeed, several reports have identified early life critical periods for microbiota influences on behavioral and neurochemical endpoints in germ free mice ^5,76^. However, the age-specific profile of sugar-associated microbiome dysbiosis and neurocognitive impairments remains to be determined.

Given that the adolescent rats consuming SSBs compensated for these calories by consuming less chow, it is possible that reduced nutrient (e.g., dietary protein) consumption may have contributed to the deficits in hippocampal function. However, we think this is unlikely, as adolescent SSB access did not produce any substantial nutrient deficiency that would restrict growth, as evidenced by the similarities in body weight between the experimental and control group. Furthermore, prior studies that directly examined the effects of adolescent caloric (and thereby nutrient) restriction on learning and memory in rats found that there were no differences in hippocampal-dependent memory function when rats were restricted by ∼40% from PN 25-PN 67 ^68^, Importantly, the parameters in this study closely match those in the present study, as our adolescent SSB access was given over a similar developmental period prior to behavioral testing, and produced a ∼40% reduction in total chow kcal consumption. Thus, it is likely that excessive sugar consumption and not nutrient deficiency led to the memory deficits, although future work is needed to more carefully examine these variables independently.

While our study reveals a strong negative correlation between levels of fecal *Parabacteroides* and performance in the hippocampal-dependent contextual episodic memory NOIC task, as well as impaired NOIC performance in rats given access to a sugar solution during adolescence, sugar intake did not produce impairments in the perirhinal cortex-dependent NOR memory task. This is consistent with our previous report in which rats given access to an 11% sugar solution during adolescence were impaired in hippocampal-dependent spatial memory (Barne’s maze procedure), yet were not impaired in a nonspatial task of comparable difficulty that was not hippocampal-dependent ^24^. Present results revealing that early life sugar consumption negatively impacts hippocampal-dependent contextual-based object recognition memory (NOIC) without influencing NOR memory performance is also consistent with previous reports using a cafeteria diet high in both fat content and sugar ^69,70^. On the other hand, enrichment of *P. johnsonii* and *P. distasonis* in the present study impaired memory performance in both tasks, suggesting a broader impact on neurocognitive functioning with this targeted bacterial enrichment approach.

Gene pathway analyses from dorsal hippocampus RNA sequencing identified multiple neurobiological pathways that may functionally connect gut dysbiosis with memory impairment. Early life sugar consumption was associated with alterations in several neurotransmitter synaptic signaling pathways (e.g., glutamatergic, cholinergic) and intracellular signaling targets (e.g., cAMP, MAPK). A different profile was observed in *Parabacteroides*-enriched animals, where gene pathways involved with metabolic function (e.g., fatty acid oxidation, branched chain amino acid degradation) and neurodegenerative disease (e.g., Alzheimer’s disease) were altered relative to controls. Given that sugar has effects on bacterial populations in addition to *Parabacteroides*, and that sugar consumption and *Parabacteroides* treatment differentially influenced peripheral glucose metabolism and body weight, these transcriptome differences in the hippocampus are not surprising. However, gene clusters involved with dopaminergic synaptic signaling were significantly influenced by both early life sugar consumption and *Parabacteroides* treatment, thus identifying a common pathway through which both diet-induced and gut bacterial infusion-based elevations in *Parabacteroides* may influence neurocognitive development. Though differentially expressed genes were commonly affected in opposite directions in *Parabacteroides* enriched animals compared with early life sugar treated animals, it is possible that perturbations to the dopamine system play a role in the observed cognitive dysfunction. For example, while dopamine signaling in the hippocampus has not traditionally been investigated for mediating memory processes, several recent reports have identified a role for dopamine inputs from the locus coeruleus in regulating hippocampal-dependent memory and neuronal activity ^71,72^. Interestingly, endogenous dopamine signaling in the hippocampus has recently been linked with regulating food intake and food-associated contextual learning ^73^, suggesting that dietary effects on gut microbiota may also impact feeding behavior and energy balance-relevant cognitive processes.

It is important to note that comparisons between the gene expressional analyses in the *Parabacteroides* enrichment and sugar consumption experiments should be made cautiously given that there were slight differences in timing of the hippocampus tissue harvest between the two experiments (PN 65 for sugar consumption vs PN 83 for the *Parabacteroides* enrichment). Further, future work is needed to determine whether differences in gene expression observed in each experiment translates to differential expression at the protein level. It is also worth emphasizing that the levels of *Parabacteroides* conferred by our enrichment study were substantially higher than in the dietary sugar study, and thus it is not surprising that *Parabacteroides* enrichment would confer a different impact on host physiology, hippocampal gene expression, and neurocognition compared to *Parabacteroides* elevations associated with SSB consumption. Regardless of these caveats in comparing the two models, our data extend the field by highlighting a specific bacterial population that 1) is capable of negatively impacting neurocognitive development when experimentally enriched, and 2) is elevated by early-life consumption of dietary sugar with levels correlating negatively with hippocampal-dependent memory performance.

Many of the genes that were differentially upregulated in the hippocampus by *Parabacteroides* enrichment were involved in fat metabolism and transport. Thus, it is possible that *Parabacteroides* conferred an adaptation in the brain, shifting fuel preference away from carbohydrate toward lipid-derived ketones. Consistent with this framework, *Parabacteroides* was previously shown to be upregulated by a ketogenic diet in which carbohydrate consumption is drastically depleted and fat is used as a primary fuel source due. Furthermore, enrichment of *Parabacteroides merdae* together with *Akkermansia muciniphila* was protective against seizures in mice ^29^. It is possible that *P. distasonis* reduces glucose uptake from the gut, enhances glucose clearing from the blood, and/or alters nutrient utilization in general, an idea further supported by recent finding that *P. distasonis* is associated with reduced diet- and genetic-induced obesity and hyperglycemia in mice ^48^.

The present findings produce several opportunities for further mechanistic investigation. For example, how do diet-induced alterations in gut bacteria impact the brain? Several possible mechanisms have been investigated and proposed, such as impaired gut barrier function and endotoxemia ^64,74^, perhaps related to altered short chain fatty acid production ^67,75^. Moreover it is well known that the liver is negatively impacted by excessive fructose consumption ^76^, and emerging evidence highlights a gut microbiome-liver axis with crosstalk via bile acids and cytokines ^77^. It is possible that dietary sugar induced microbiota changes alter the hepatic-gut axis, thus contributing to altered cognitive function. Indeed, an altered bile acid profile due to gut microbiota produced bile acid secondary metabolites is associated with cognitive dysfunction in Alzheimer’s Disease in humans ^78^.

Taken together, our collective results provide insight into the neurobiological mechanisms that link early life unhealthy dietary patters with altered gut microbiota changes and neurocognitive impairments. Currently probiotics, live microorganisms intended to confer health benefits, are not regulated with the same rigor as pharmaceuticals but instead are sold as dietary supplements. Our findings suggest that gut enrichment with certain species of *Parabacteroides* is potentially harmful for neurocognitive mnemonic development. These results highlight the importance of conducting rigorous basic science analyses on the relationship between diet, microorganisms, brain, and behavior prior to widespread recommendations of bacterial microbiome interventions for humans.

## Acknowledgements

We thank Alyssa Cortella for contributing the rodent artwork. We thank Caroline Szjewski, Lekha Chirala, Vaibhav Konanur, Sarah Terrill, and Ted Hsu for their critical contributions to the research. The research was supported by DK116942, DK104897, and institutional funds to S.E.K., DK118000 and DK111158 to E.E.N., DK116558 to A.N.S., DK 118944 to C.M.L. C.A.O. was supported by an F31 AG064844. E.Y.H. was supported by ARO MURI award W911NF-17-1-0402. DK104363 to X.Y., Eureka Scholarship and BWF-CHIP Fellowship to Y.C.

---

**Figure S1:**
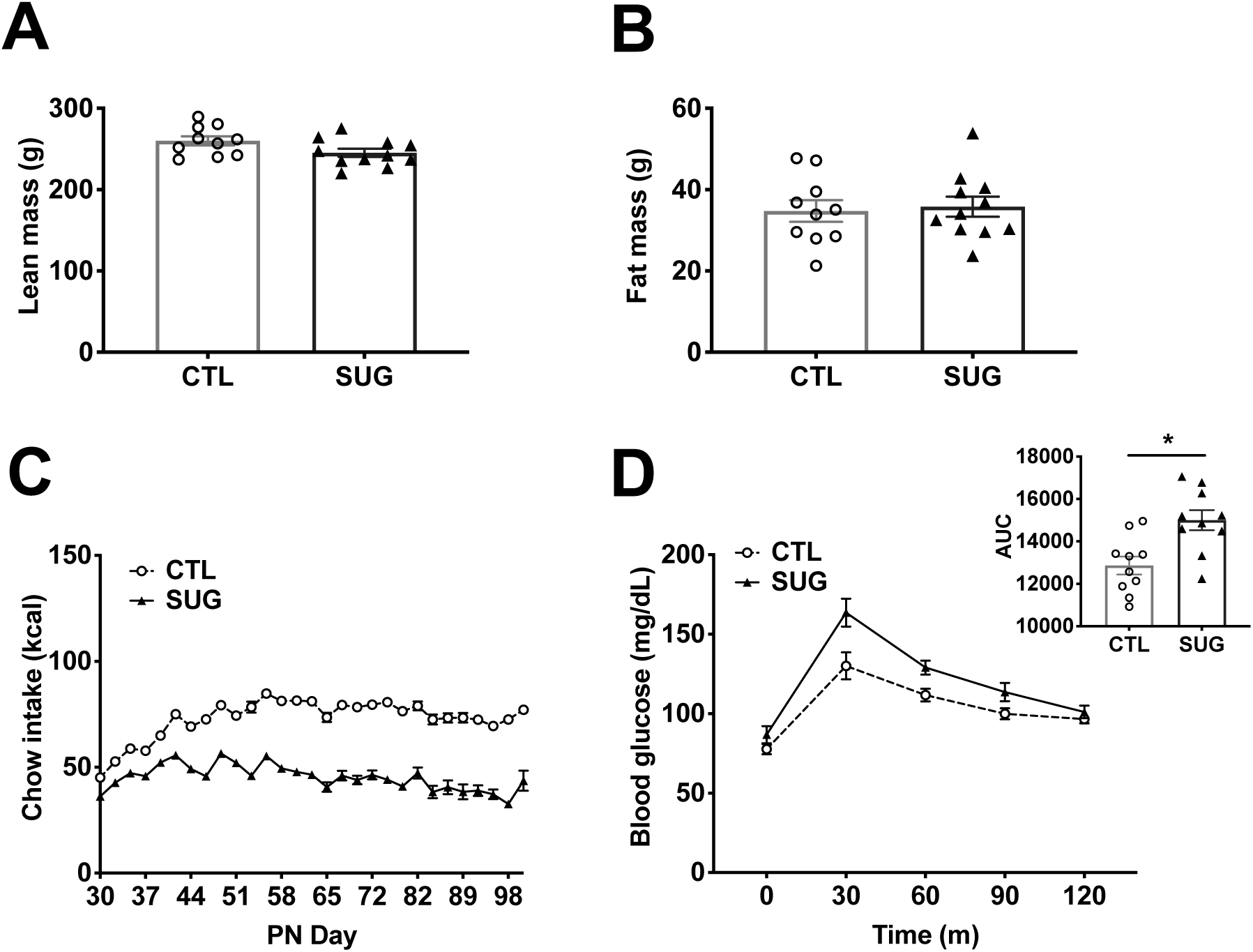
Effect of early life sugar consumption on food intake and metabolic measures. (A, B) There were no differences in lean mass or in fat mass between animals fed sugar solutions or control animals (n=10,11; two-tailed, type 2 Student’s T-test). (C) Kcals from chow intake were lower throughout the feeding period in animals fed early life sugar (n=10,11). (D) Results from the intraperitoneal glucose tolerance test show an elevated area under the curve (AUC) in rodents fed sugar solutions during early life (n=10,11; two-tailed, type 2 Student’s T-test; P<.05). CTL=control, SUG= sugar, PN= post-natal day; data shown as mean ± SEM.

**Figure S2:**
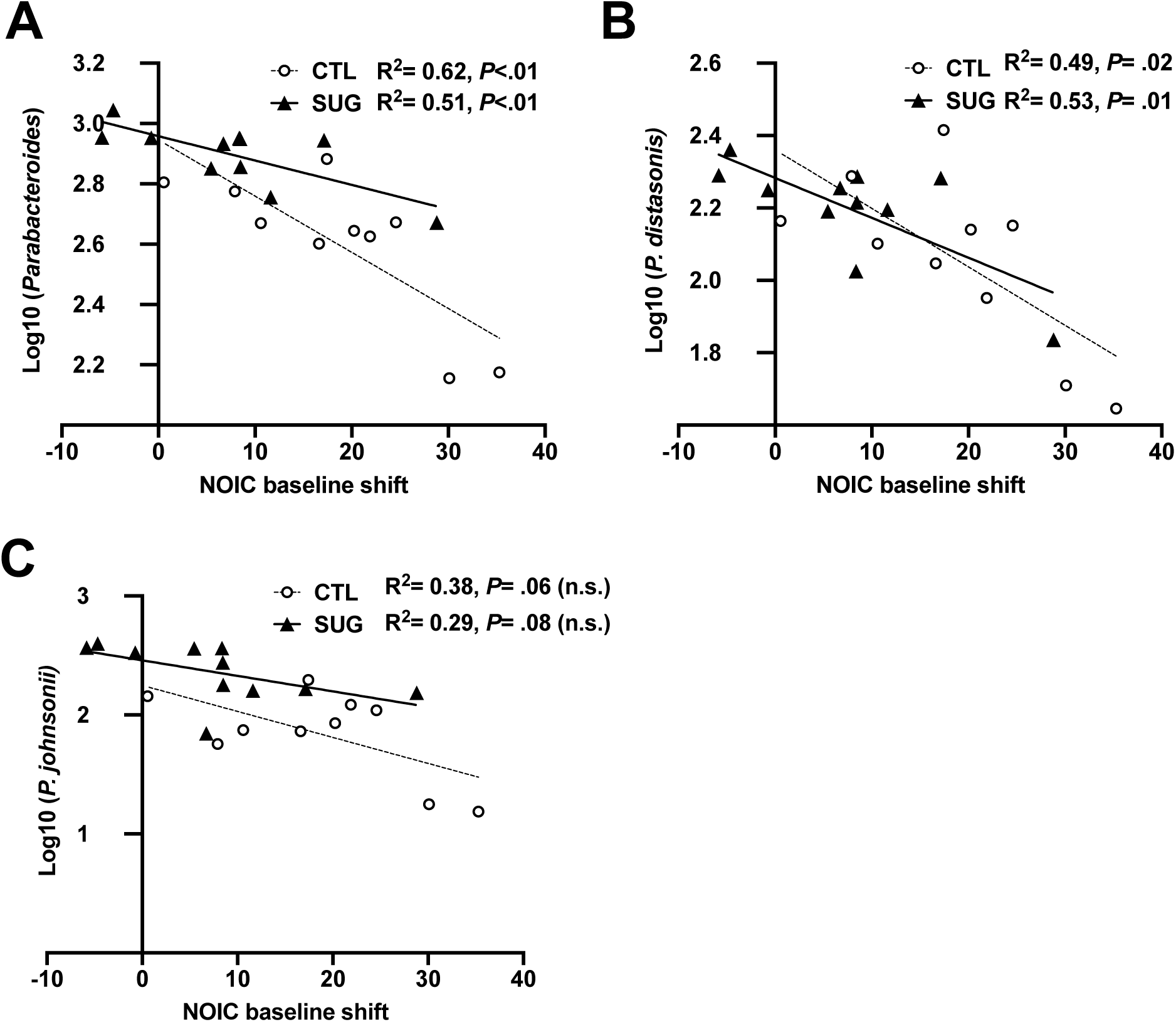
Relationship between *Parabacteroides* and behavioral outcomes in the Novel Object in Context task (NOIC) A) Linear regression of log normalized fecal *Parabacteroides* counts against shift from baseline performance scores in the NOIC task in sugar (SUG) and control (CTL) groups (n=10, 11). (B, C) Linear regression of the most abundant fecal *Parabacteroides* species against shift from baseline performance scores in NOIC across all groups tested (n=10, 11). *P<0.05; data shown as mean ± SEM.

**Figure S3:**
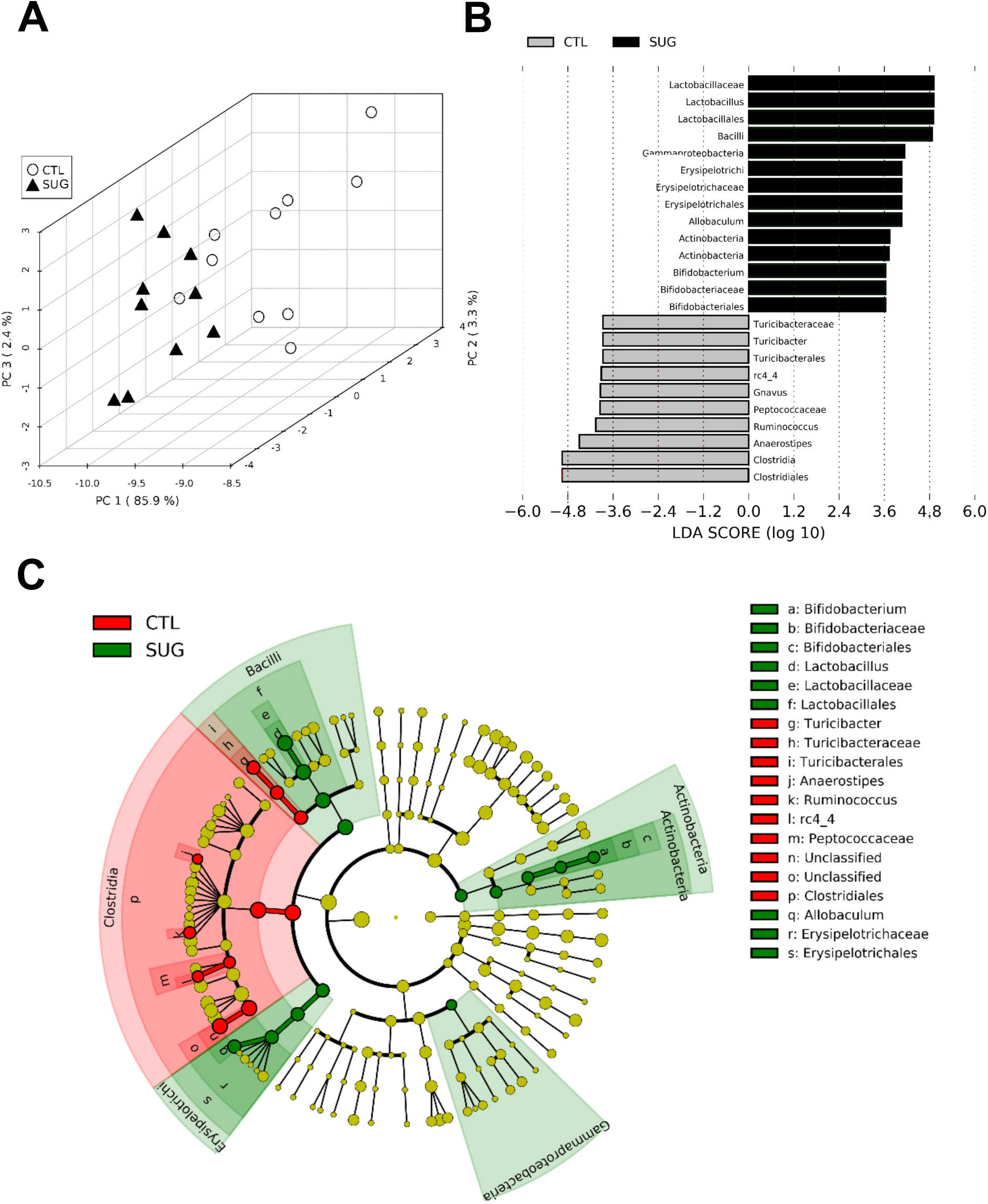
Effect of early life sugar consumption on the rat cecal microbiota. (A) Principal component analysis (PCA) was run using all phylogenic levels (112 normalized taxa abundances) and shows different clustering patterns based on overall cecal microbial profiles. (B) Linear discriminant analysis (LDA) Effect Size (LEfSe), run using the GALAXY platform, identified characteristic features of the cecal microbiota of rats fed a control diet or early life sugar. Relative differences among groups were used to rank the features with the LDA score set at 2. (C) Identified taxa are displayed by scores and on a phylogenic cladogram. CTL=control, SUG= sugar.

**Figure S4:**
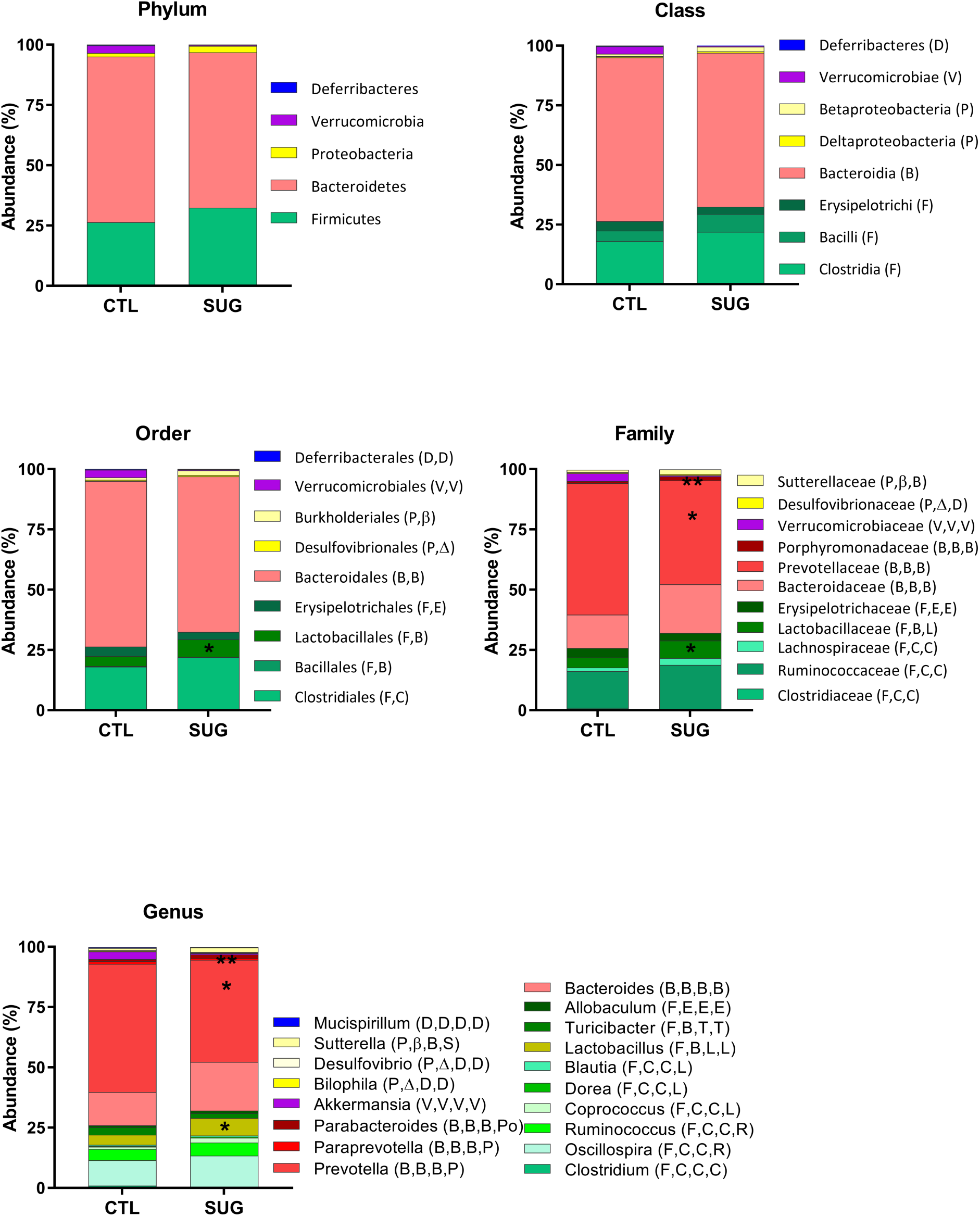
Effect of early life sugar consumption on the rat fecal microbiota. Filtered bacterial abundances by taxonomic levels phylum, class, order, family, genus in fecal samples from rats fed a control diets or early life sugar. Differences in abundances were assessed by Mann-Whitney non-parametric test. * p<0.05, *** p<0.001. CTL=control, SUG= sugar.

**Figure S5:**
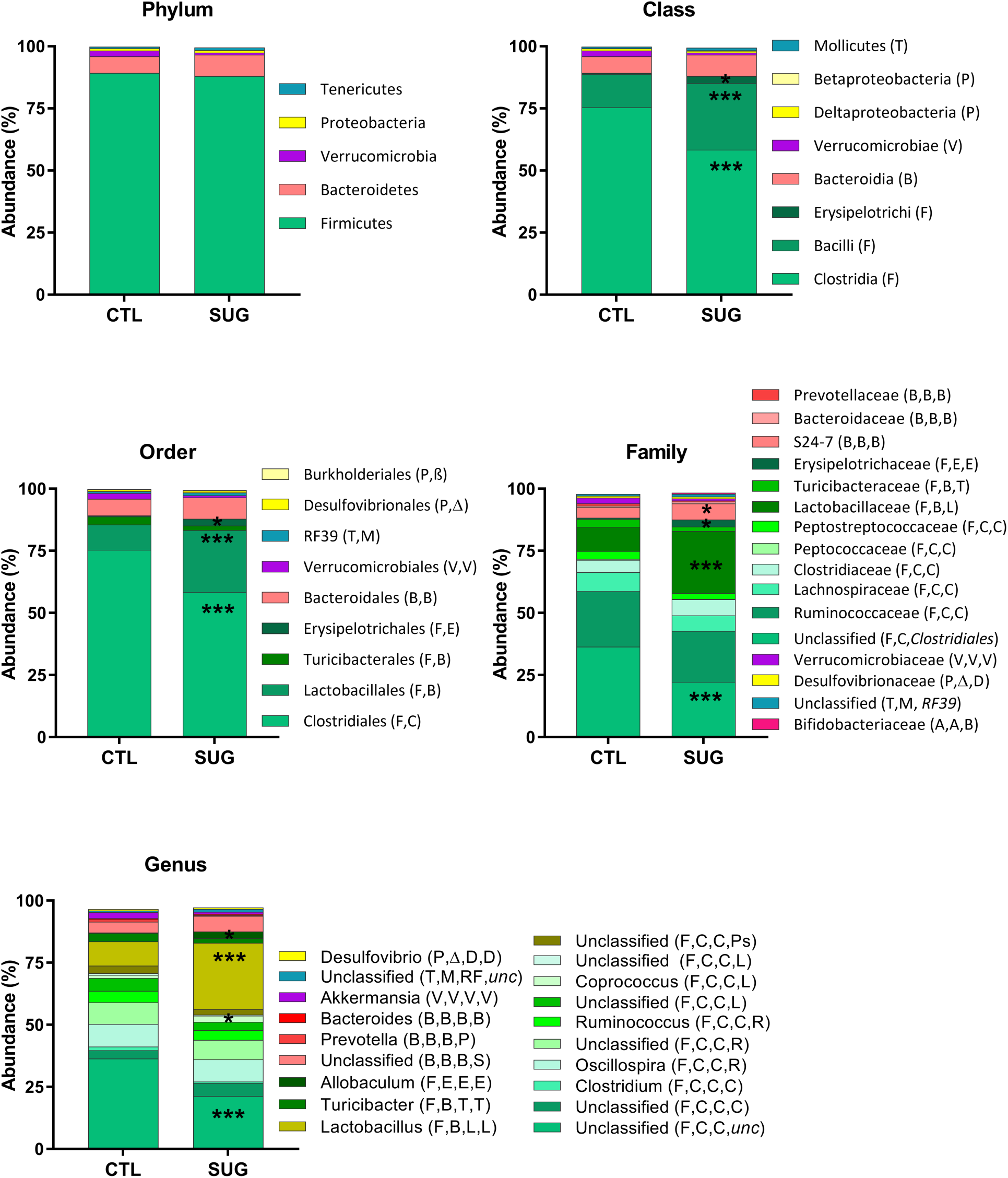
Effect of early life sugar consumption on the rat cecal microbiota: Filtered bacterial abundances by taxonomic levels phylum, class, order, family, genus in cecal samples from rats fed a control diets or early life sugar. Differences in abundances were assessed by Mann-Whitney non-parametric test. * p<0.05, *** p<0.001. CTL=control, SUG= sugar.

**Phylogenic taxonomy legend for Figure S4, S5**:

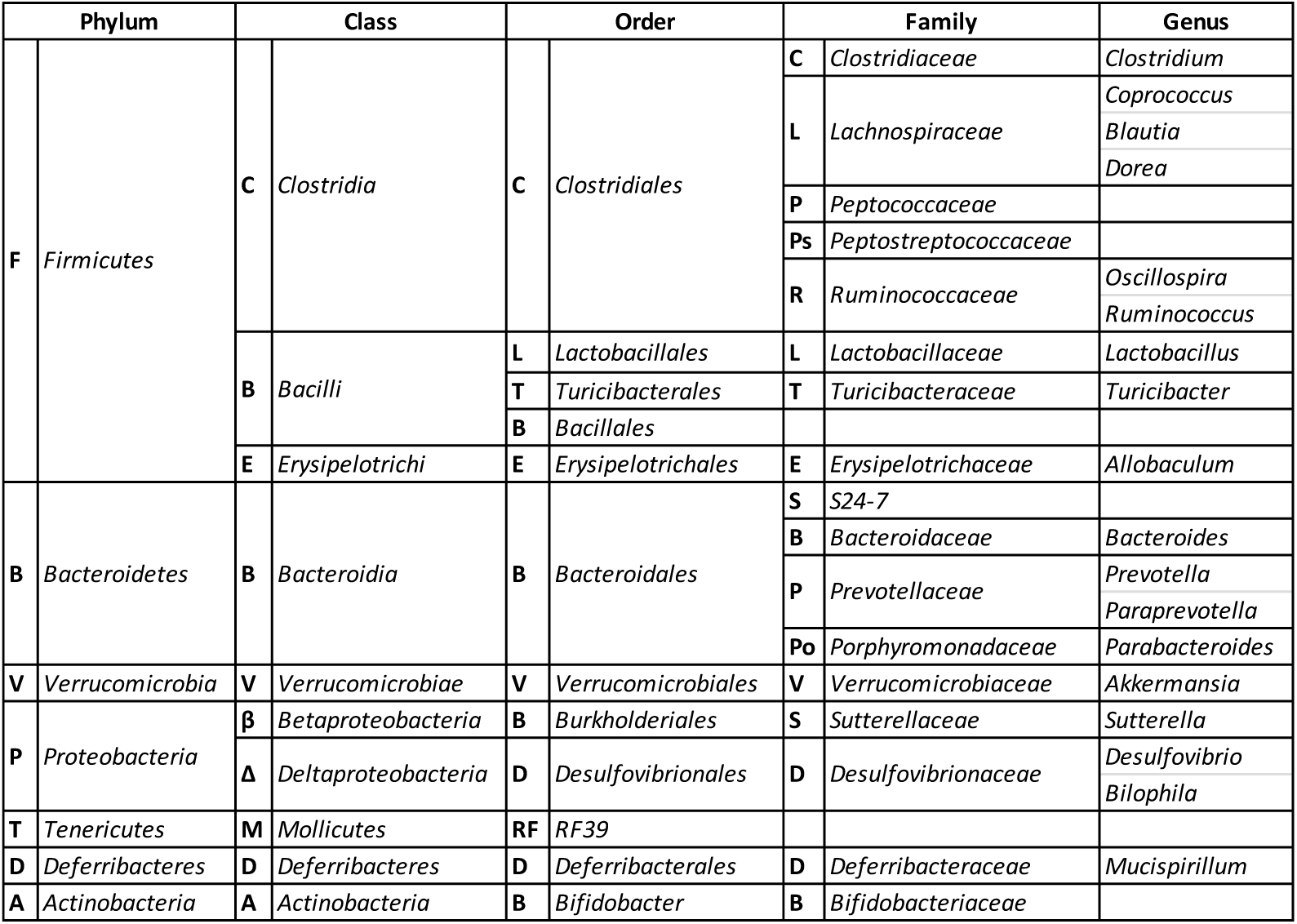

**Figure S6:**
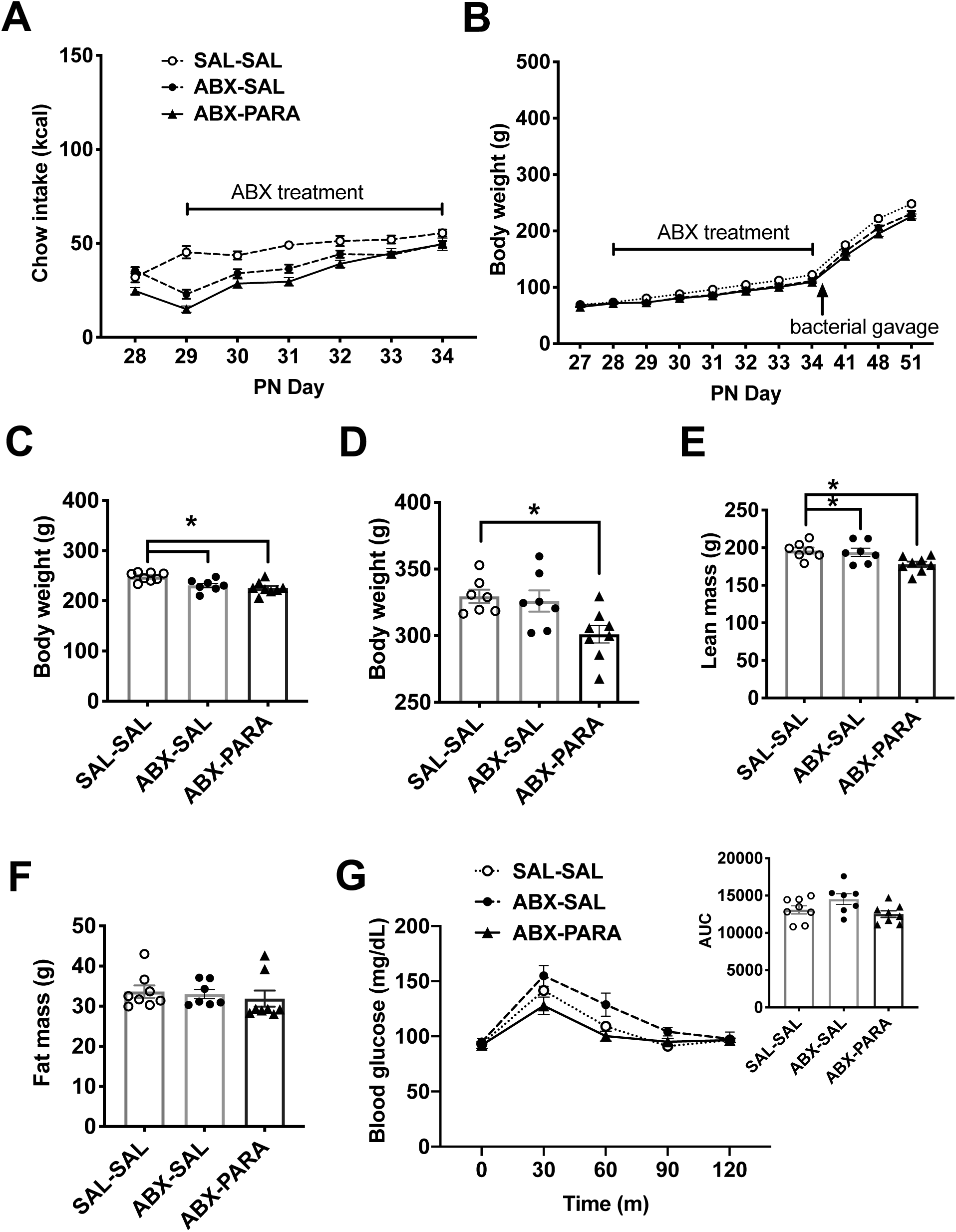
Experimental design, food intake, and metabolic measures for gut *Parabacteroides* enrichment. (A) Effect of antibiotic treatment on food intake and body weight (n=7,8). (C) Effect of gut *Parabacteroides* enrichment on body weight at PN 51 prior to the start of behavioral testing (n=7,8; one way ANOVA with Tukey’s post hoc test, F_(2,20)_= 8.79; *P<.05) (D) Effect of gut *Parabacteroides* enrichment on body weight (n=7,8; one way ANOVA with Tukey’s post hoc test, F_(2,19)_= 5.7; *P<.05) (E) lean mass (n=7,8; one way ANOVA with Tukey’s post hoc test, F_(2,19)_= 5.33; *P<.05) (F) and body fat (one way ANOVA, n.s.) at PN 76. (G) Blood glucose levels during an interaperitoneal glucose tolerance test (IP GTT) (n=7,8 one way ANOVA for AUC; n.s.) SAL-SAL=saline-saline control, ABX-SAL= antibiotics-saline control, ABX-PARA= antibiotics-*P. johnsonii* and *P. distasonis* enriched, PN= post-natal day; data shown as mean ± SEM.

**Figure S7:**
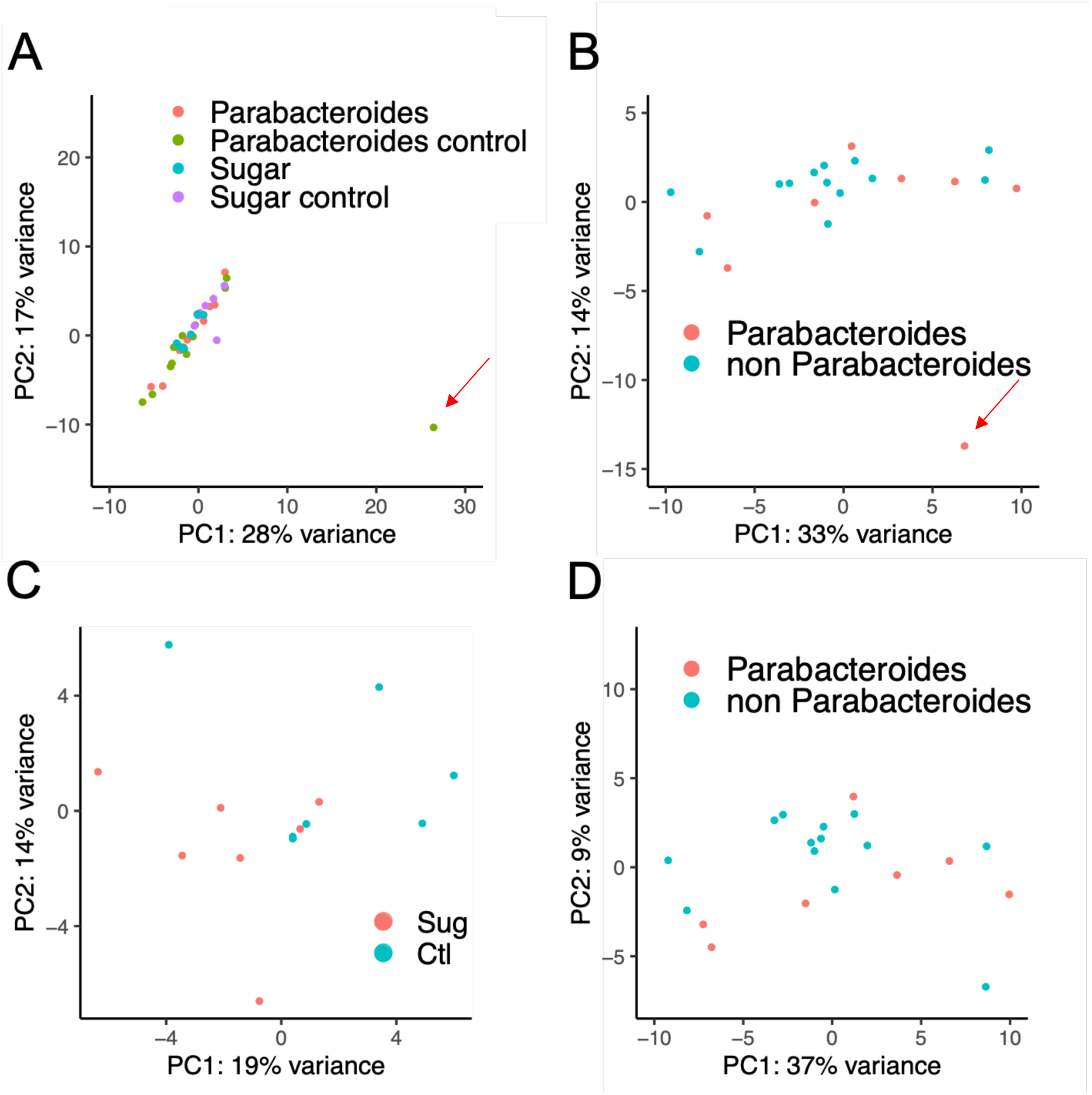
Principal component analyses (PCA) of hippocampal gene expression data to identify outliers. (A) PCA identified one control sample (red arrow) as an outlier when all samples from both sugar and *Parabacteroides* enrichment experiments were considered. (B) PCA identified one treatment sample (red arrow) from the *Parabacteroides* experiment as an outlier. After removing the outliers, PCA for the remaining samples from the sugar treatment experiment (C) and those from the *Parabacteroides* enrichment experiments (D).

**Figure S8:**
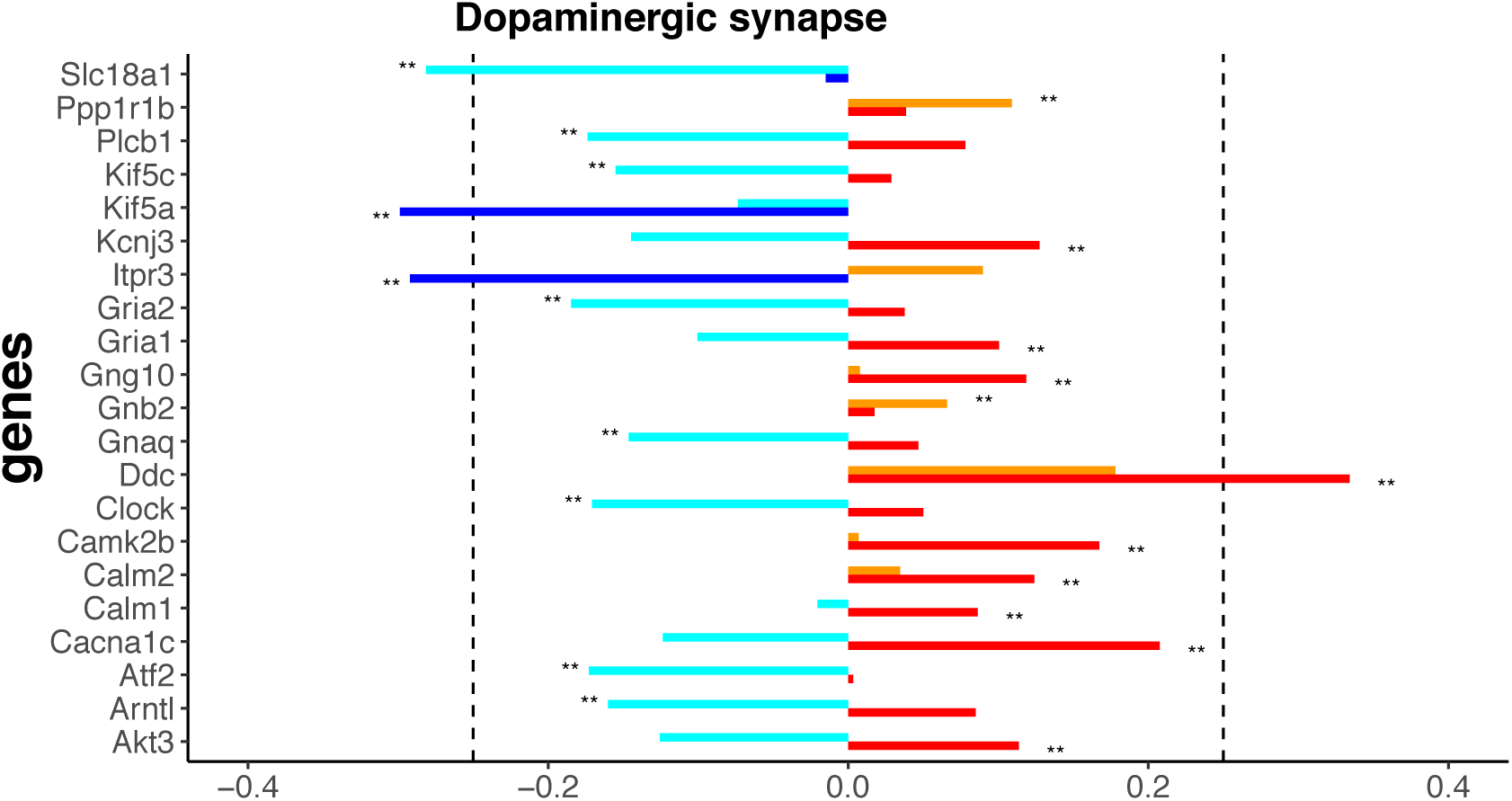
Comparison of hippocampal gene expression pathways altered by sugar and *Parabacteriodes.* The dopaminergic synapse pathway overlaps in the sugar and *Parabacteroides* transfer experiments. Red= upregulated by sugar, dark blue= downregulated by sugar, orange= upregulated by *Parabacteroides*, light blue= downregulated by *Parabacteroides.* * *P* < 0.05 and ** *P* < 0.01. Dotted line indicated ± 0.25 log2 fold change.

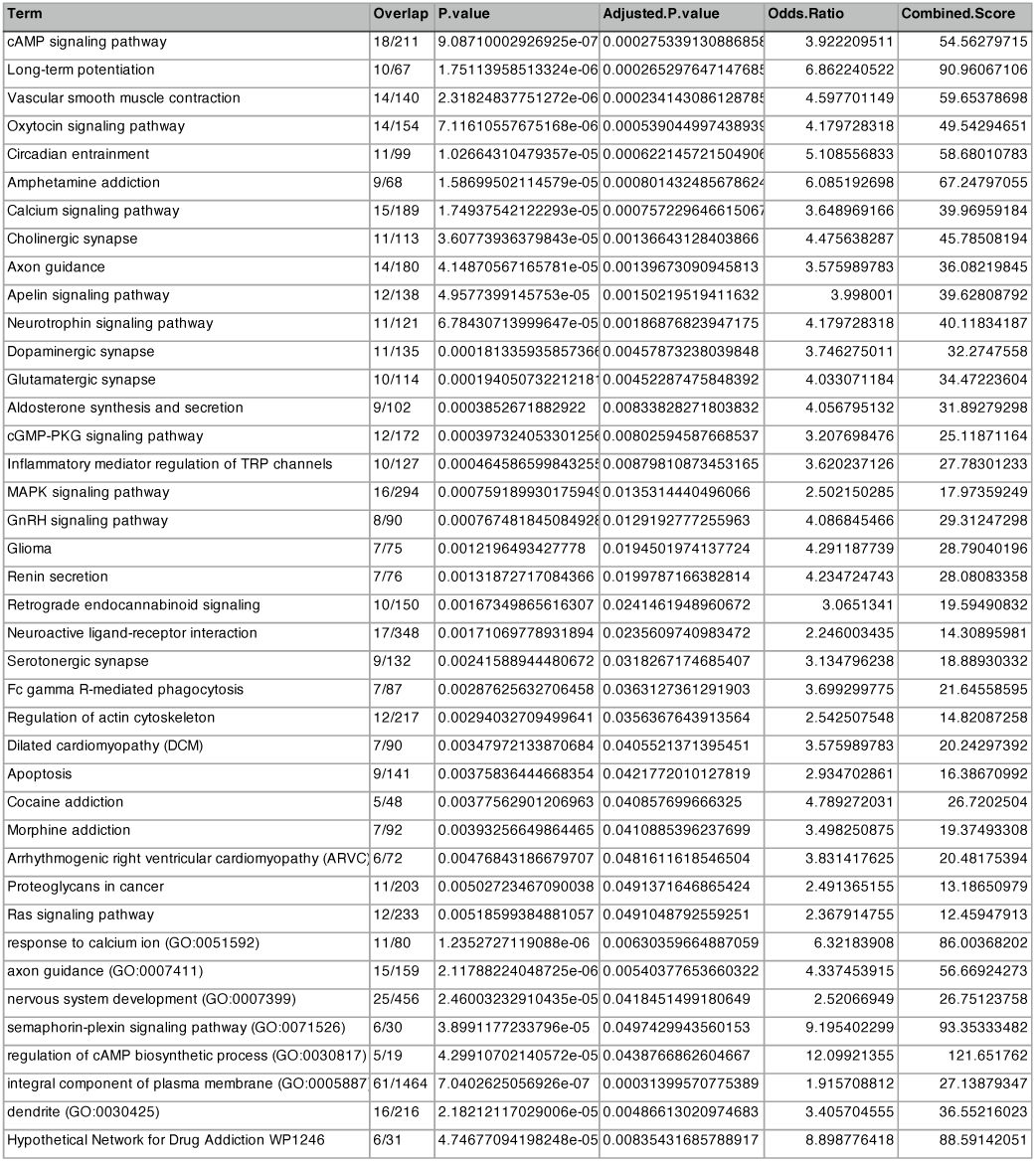

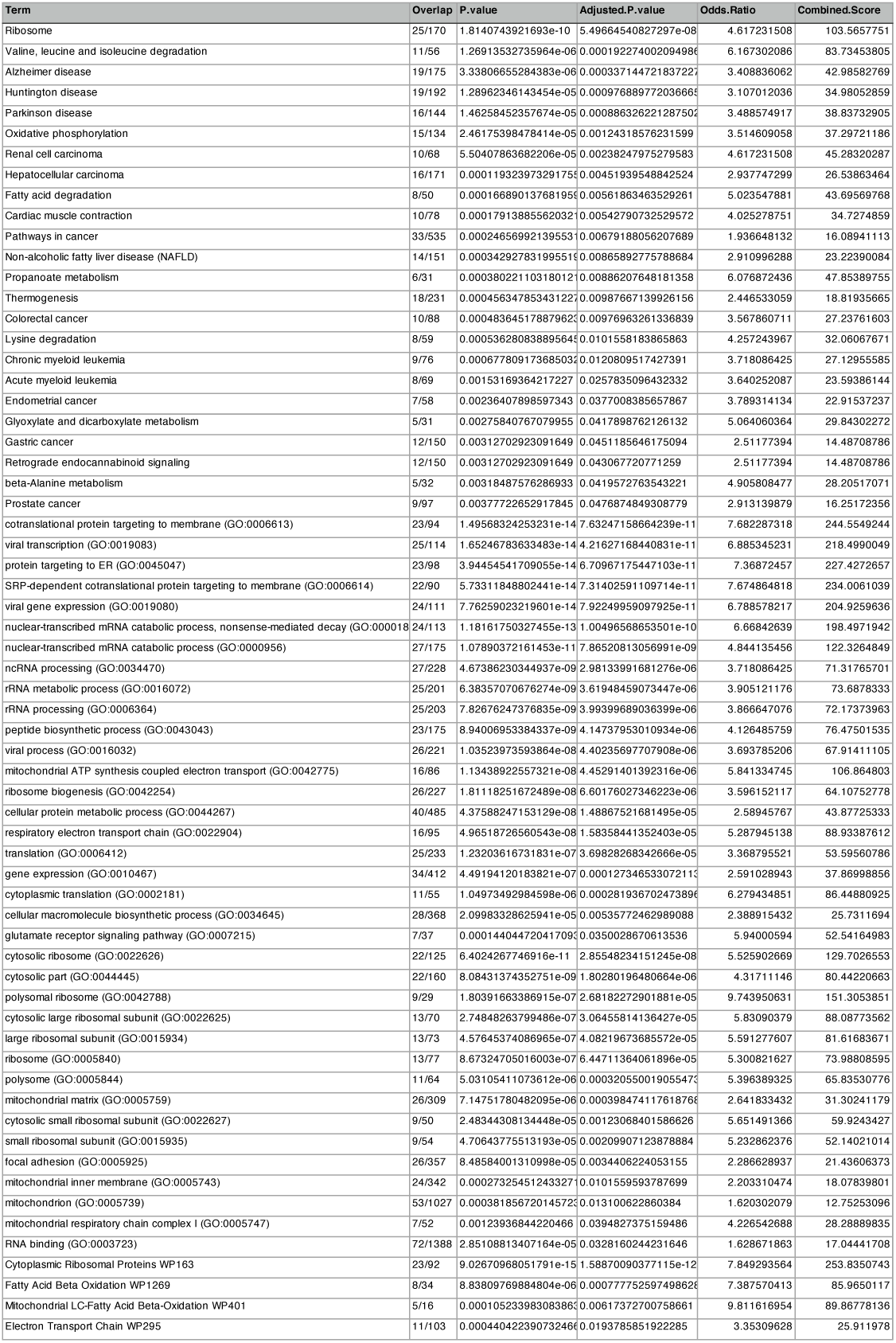

